# The genotype-phenotype landscape of an allosteric protein

**DOI:** 10.1101/2020.09.30.320812

**Authors:** Drew S. Tack, Peter D. Tonner, Abe Pressman, Nathanael D. Olson, Sasha F. Levy, Eugenia F. Romantseva, Nina Alperovich, Olga Vasilyeva, David Ross

**Affiliations:** National Institute of Standards and Technology, Gaithersburg, MD, 20899, USA; SLAC National Accelerator Laboratory, Menlo Park, CA, 94025, USA; Joint Initiative for Metrology in Biology, Stanford, CA, 94305, USA

## Abstract

Allostery is a fundamental biophysical mechanism that underlies cellular sensing, signaling, and metabolism. Yet a quantitative understanding of allosteric genotype-phenotype relationships remains elusive. Here we report the large-scale measurement of the genotype-phenotype landscape for an allosteric protein: the *lac* repressor from *Escherichia coli*, LacI. Using a method that combines long-read and short-read DNA sequencing, we quantitatively measure the dose-response curves for nearly 10^5^ variants of the LacI genetic sensor. The resulting data provide a quantitative map of the effect of amino acid substitutions on LacI allostery and reveal systematic sequence-structure-function relationships. We find that in many cases, allosteric phenotypes can be quantitatively predicted with additive or neural-network models, but unpredictable changes also occur. For example, we were surprised to discover a new band-stop phenotype that challenges conventional models of allostery and that emerges from combinations of nearly silent amino acid substitutions.

## Introduction

Allostery is a fundamental biophysical mechanism that underlies cellular regulatory processes including sensing, signaling, and metabolism^1–3^. With allosteric regulation, ligand binding at one site on a biomolecule causes a conformational change that affects the activity of another, often distal, site. This conformational switching provides a sense-and-response function that defines the allosteric phenotype. Quantitative descriptions relating that phenotype to its causal genotype would improve our understanding of cellular function and evolution, and advance protein design and engineering^4–6^. However, the intramolecular interactions that mediate allosteric regulation are complex and distributed widely across the biomolecular structure, making the development of general quantitative descriptions challenging.

Recently described fitness landscape approaches have enabled the phenotypic characterization of 10^4^ to 10^5^ genotypes simultaneously^7–12^. Measurements at this scale facilitate the exploration of genotypes with widely distributed mutations, making them ideal for probing complex biological mechanisms like allostery. However, to quantitatively characterize the sense-and-response phenotypes inherent to allostery, a measurement must encompass the full dose-response curve that describes biomolecular activity as a function of ligand concentration.

Genetic sensors have served as a model of allosteric regulation for decades, and today are central to engineering biology. Genetic sensors are allosteric proteins that regulate gene expression in response to stimuli, giving cells the ability to regulate their metabolism and respond to environmental changes. Like many allosteric genetic sensors, the *lac* repressor, LacI, binds to DNA upstream of regulated genes, preventing transcription. Ligand binding to LacI causes a switch from the DNA-binding conformation to a non-binding conformation that allows transcription to proceed. This conformational switching results in the allosteric phenotype that is quantitatively defined by a dose-response curve relating the concentration of input ligand (*L*) to the output response (the expression level of regulated genes, *G*). Genetic sensors typically have sigmoidal dose-response curves following the Hill equation:

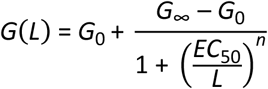

where *G*_0_ is basal gene expression in the absence of ligand, *G*_∞_ is gene expression at saturating ligand concentrations, *EC*_50_ is the effective concentration of ligand that results in gene expression midway between *G*_0_ and *G*_∞_, and *n* quantifies the steepness of the dose-response curve (Fig. 1e). The dose-response curve is affected by several biophysical constants including ligand-binding affinity, DNA-binding affinity, and the allosteric constant (the equilibrium between the two conformations)^2,13–15^. These constants depend on amino acid residues and interactions spread widely across the protein, making it difficult to predict the effects of changes in protein sequence.

**Fig. 1.**
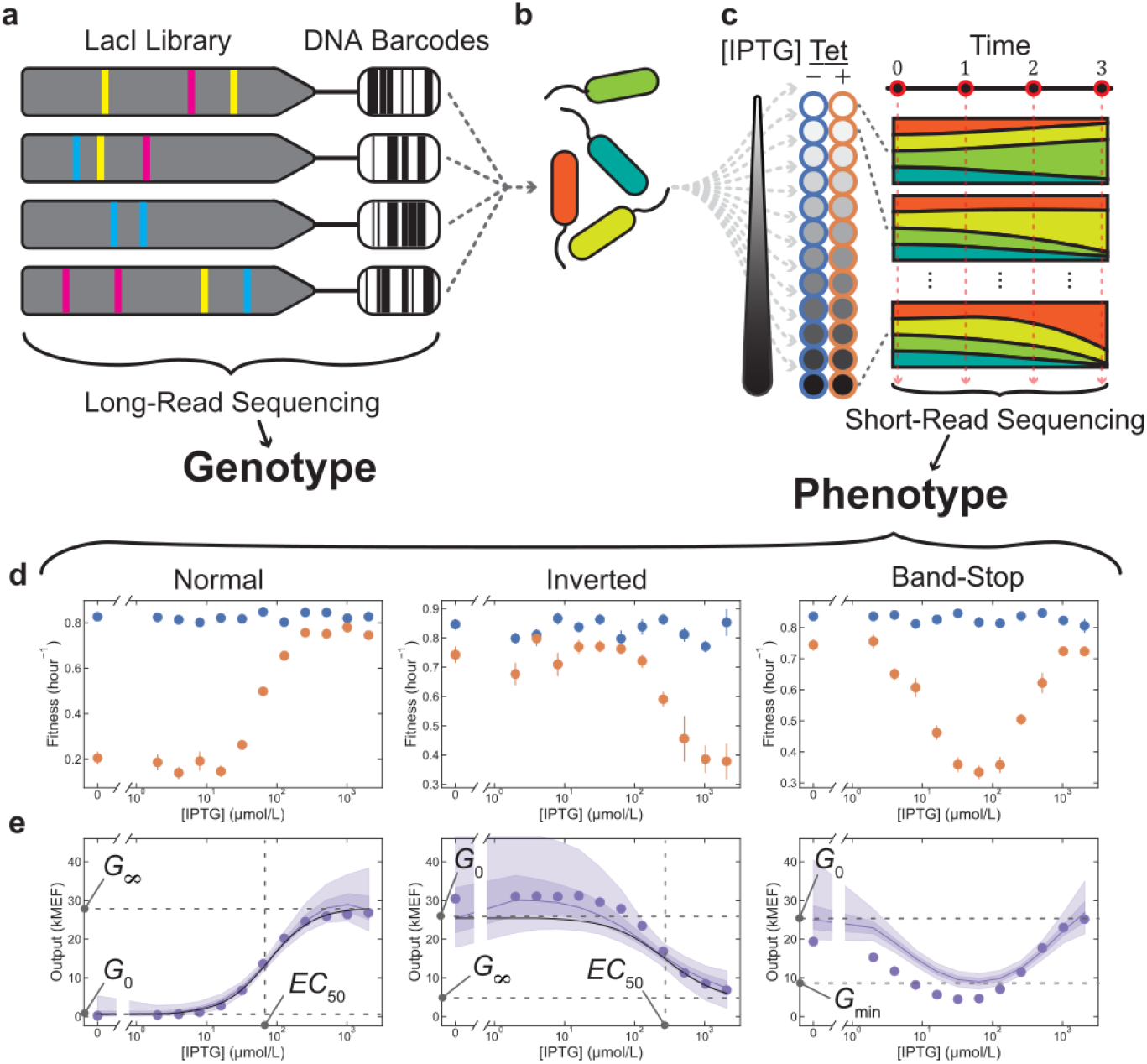
Large-scale allosteric genotype-phenotype landscape measurement. **a** A library of *lac* repressor (LacI) variants was generated by random mutagenesis of the *lacI* coding DNA sequence (CDS). The CDS for each variant was attached to a DNA barcode and inserted into a plasmid where the LacI variant regulated expression of a tetracycline resistance gene. The CDS and corresponding barcode on each plasmid were determined with long-read sequencing. **b** The library was transformed into *E. coli*. **c** Cells containing the library were grown in 24 chemical environments, including 12 concentrations of the ligand IPTG, each with (orange) and without (blue) tetracycline. Cultures were maintained in exponential growth. Changes in the relative abundance of each variant were measured with short-read sequencing of DNA barcodes at four timepoints and were used to determine the fitness associated with each variant in each environment. **d** The fitness without tetracycline (blue) is independent of IPTG concentration. The fitness with tetracycline (orange) depends on the IPTG concentration via the dose-response of each variant. **e** Dose-response curves for 62,472 LacI variants were determined from the fitness measurements with Bayesian inference using a Hill equation model (black lines for variants with normal and inverted dose-response curves) and a Gaussian process (GP) model (purple lines, shaded regions indicate 50% and 90% credible intervals). Flow cytometry verification measurements (purple points) generally agreed with Bayesian inference results and verified the existence of the band-stop and other phenotypes. Dose-response output was calibrated from fitness to fluorescent gene expression (Supplementary Fig. 16) and reported in molecules of equivalent fluorophore (MEF). Error bars indicate ± one standard deviation and are often within markers.

## Results

### Measuring the genotype-phenotype landscape

To measure the genotype-phenotype landscape for the allosteric LacI sensor, we first created a library of LacI variants using mutagenic PCR and attached a DNA barcode to the coding DNA sequence of each variant (Fig. 1a). We inserted the barcoded library into a plasmid where LacI regulates the expression of a tetracycline resistance gene (Supplementary Fig. 1a). Consequently, in the presence of tetracycline, the LacI dose-response modulates cellular fitness based on the concentration of the input ligand isopropyl-β-D-thiogalactoside (IPTG). We then transformed the library into *E. coli* for the landscape measurement (Fig. 1b). To ensure that most variants in the library could regulate gene expression, we used fluorescence-activated cell sorting (FACS) to enrich the library for variants with low *G*_0_. Then, using high-accuracy, long-read sequencing^16^, we determined the genotype for every variant in the library and indexed each variant to its attached DNA barcode (Fig. 1a).

The library contained 62,472 different LacI genotypes, with an average of 7.0 single nucleotide polymorphisms (SNPs) per genotype. Many SNPs were synonymous, i.e. coded for the same amino acid, so the library encoded 60,398 different amino acid sequences with an average of 4.4 amino acid substitutions per variant (Supplementary Fig. 2b).

**Fig. 2.**
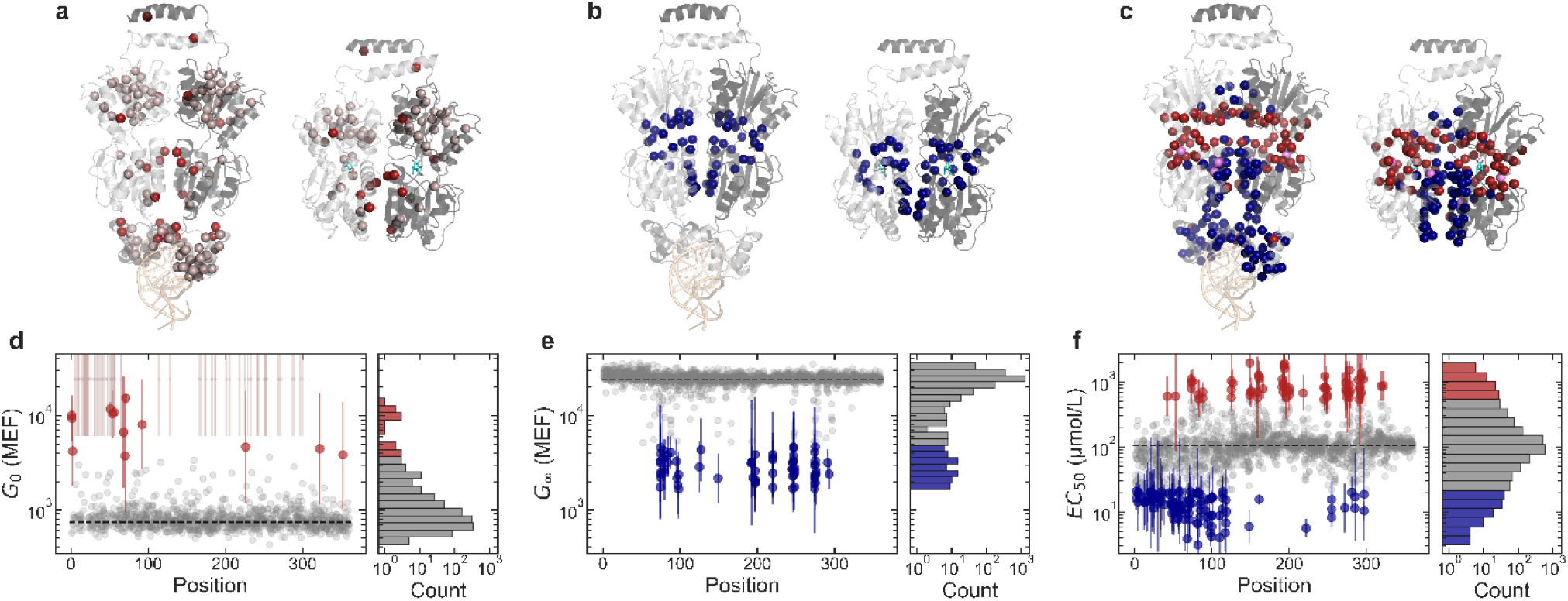
Effect of single amino acid substitutions on allosteric function. **a-c** Protein structures showing the locations of amino acid substitutions that affect each Hill equation parameter: *G*_0_ (**a**), *G*_∞_ (**b**), *EC*_50_ (**c**). For each, the DNA-binding configuration is shown on the left (DNA in light orange, PDB ID: 1LBG^22^) and the ligand-binding configuration is shown on the right (IPTG in cyan, PDB ID: 1LBH^22^). Both configurations are shown with the view oriented along the protein dimer interface, with one monomer in light gray and the other monomer in dark gray. Colored spheres highlight residues where substitutions cause a greater than 5-fold change in the Hill equation parameter relative to wild-type LacI. Red spheres indicate residues where substitutions increase the parameter, and blue spheres indicate residues where substitutions decrease the parameter. At three residues (A82, I83, and F161), some substitutions decrease *EC*_50_, while other substitutions increase *EC*_50_ (violet spheres in **c**). **d-f** Scatter plots show the effect of each substitutions as a function of position. Substitutions that change the parameter by less than 5-fold are shown as gray points. Substitutions that change the parameter by more than 5-fold are shown as red or blue points with error bars. In **a** and **d**, gray-pink spheres and points indicate positions for substitutions that are completely missing from the landscape dataset and that have been shown by previous work to result in constitutively high *G*(*L*)^18,19^. Histograms to the right of each scatter plot show the overall distribution of single-substitution effects. *G*_0_ and *G*_∞_ are reported in molecules of equivalent fluorophore (MEF). Error bars indicate ± one standard deviation.

To quantitatively determine the allosteric phenotype for every LacI variant in the library, we developed a new method to characterize the dose-response curves for large genetic sensor libraries. Briefly, we grew *E. coli* containing the library in 24 chemical environments (12 ligand concentrations, each with and without tetracycline). We used short-read sequencing of the DNA barcodes to measure the relative abundance of each variant at four timepoints during growth (Fig. 1c). We then used the changes in relative abundance to determine the fitness associated with each variant in each environment (Fig. 1d). Finally, for each variant in the library, we used the fitness difference (with vs. without tetracycline) from all 12 ligand concentrations to quantitatively determine the dose-response curve using Bayesian inference (Fig. 1e). Most variants had sigmoidal dose-response curves (e.g. Supplementary Figs. 3-4), which we quantitively described using the parameters of the Hill equation. We compared the results of variants with synonymous coding sequences and found that synonymous SNPs did not measurably impact the dose-response. So, for subsequent analysis we considered only amino acid substitutions.

**Fig. 3.**
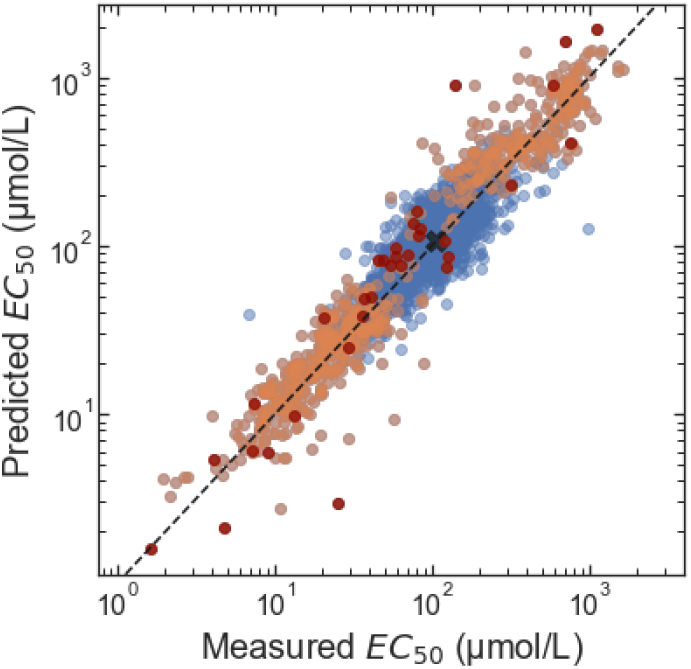
The effects of amino acid substitutions on *EC*_50_ are log-additive. The predicted *EC*_50_ for double-substitution LacI variants (i.e. two amino acid substitutions) was calculated assuming log-additivity of the effect of each single substitution on the *EC*_50_ relative to the wild-type: (*EC*_50,*AB*_ – *EC*_50,*wt*_) = (*EC*_50,*A*_ – *EC*_50,*wt*_) + (*EC*_50,*B*_ – *EC*_50,*wt*_), where ‘*wt*’ indicates the wild-type, ‘*A*’ and ‘*B*’ indicate the single-substitution variants, and ‘*AB*’ indicates the double-substitution variant. The measured *EC*_50_ of double-substitution variants is from the large-scale landscape measurement. Orange points mark double-substitution variants in which one of the single substitutions causes a greater than 2.5-fold change in *EC*_50_. Dark red points mark double-substitution variants in which both single substitutions cause a greater than 2.5-fold change in *EC*_50_. The *EC*_50_ of wild-type LacI is marked with a black ‘X’. For this analysis, only experimental data was used (no results from the DNN model). Also, only data from LacI variants with low *EC*_50_ uncertainty were used (std(log_10_(*EC*_50_)) < 0.35).

To test the accuracy of the new method for library-scale dose-response curve measurements, we independently verified the results for over 100 LacI variants from the library. For each verification measurement, we chemically synthesized the coding DNA sequence for a single variant and inserted it into a plasmid where LacI regulates the expression of a fluorescent protein. We transformed the plasmid into *E. coli* and measured the resulting dose-response curve with flow cytometry (e.g. Fig. 1e). The flow cytometry results confirmed both the qualitative and quantitative accuracy of the new method (Supplementary Figs. 3-7).

### Effects of amino acid substitutions on LacI phenotype

During library construction, we chose the mutation rate to simultaneously achieve two objectives: exploration of a broad genotype-phenotype space, and acquisition of the single-amino-acid-substitution data most useful for building quantitative biophysical models of allosteric function^2,13,14^. Starting from the wild-type DNA sequence for LacI, there were 2110 possible SNP-accessible amino acid substitutions. Most of those substitutions were present in one or more variants within the library. However, nearly half were found only in combination with other substitutions. So, to comprehensively determine the impact of single amino acid substitutions, we constructed a deep neural network model (DNN) capable of accurately predicting the Hill equation parameters for LacI variants that were not directly measured. We adapted a recurrent architecture that captures the context dependence of mutational effects (Supplementary Fig. 8) and used an approximate Bayesian variational method to estimate uncertainties for the model predictions^17^.

We trained the model to predict the Hill equation parameters *G*_0_, *G*_∞_, and *EC*_50_ (Supplementary Fig. 9). For all three parameters, the root-mean-square error (RMSE) for the model predictions increases with the number of amino acid substitutions relative to the wild type (Supplementary Fig. 10). Importantly, for single-substitution variants, the model RMSE is comparable to the experimental measurement uncertainty (Supplementary Fig. 11). So, we could confidently integrate the experimental and DNN results to provide a nearly complete map of the effects of SNP-accessible amino acid substitutions. Furthermore, by integrating information about the causal substitutions from multiple genetic backgrounds, the model provided improved estimates of *EC*_50_ and *G*_∞_ for variants with *EC*_50_ near or above the maximum ligand concentration measured (Supplementary Fig. 12).

The resulting map of single-substitution effects (Supplementary Data 1) includes quantitative point estimates and uncertainties for the Hill equation parameters for 94% of the possible SNP-accessible amino acid substitutions (1991 of 2110; 964 directly from measured data, and 1027 from DNN predictions). Most of the 119 substitutions missing from the dataset were probably excluded by FACS during library preparation because they caused a substantial increase in *G*_0_. These include 83 substitutions that have been shown to result in constitutively high *G*(*L*)^18,19^. Of the 1991 substitutions included in the dataset, 38% measurably affect the dose-response curve (beyond a 95% confidence bound).

The effect of any substitution depends strongly on its location within the protein structure, indicating systematic structure-function relationships underlying allostery (Fig. 2, using structural features as defined in references^20–22^). For example, substitutions that increase the basal expression, *G*_0_, by more than 5-fold are located either in helix 4 of the DNA-binding domain, along the dimer interface, in the tetramerization helix, or at the protein start codon (Fig. 2a,d). *G*_0_ quantifies gene expression in the absence of ligand. So, apart from substitutions at the start codon that reduce the number of LacI proteins per cell^23^, these substitutions probably affect either the DNA-binding affinity, the allosteric constant, or both^14^. Interestingly, substitutions in helix 4 (R51C, Q54K, and L56M) and near the dimer interface (T68N, L71Q) that increase *G*_0_ also decrease *EC*_50_ approximately 10-fold, consistent with a change in the allosteric constant^14^ (Supplementary Fig. 13a).

Amino acid substitutions that decrease ligand-saturated expression, *G*_∞_, by more than 5-fold are all located near the ligand-binding pocket or along the dimer interface (Fig. 2b,e). Six of these substitutions also increase *EC*_50_ more than 5-fold (A75T, D88N, S193L, Q248R, D275Y, and F293Y; Supplementary Fig. 13b). Except for D88N, which is at the dimer interface, these substitutions are in the ligand-binding pocket. Substitutions near the ligand-binding pocket probably change ligand-binding affinity, though studies with targeted substitutions have shown that they can also change the allosteric constant^14^.

Amino acid substitutions that change the effective concentration, *EC*_50_, are the most numerous and are spread throughout the protein structure, with approximately 9% and 20% of all substitutions causing a greater than 5-fold or 2.5-fold shift in *EC*_50_, respectively (Fig. 2c,f). The strongest effects are from substitutions in the DNA-binding domain, ligand-binding pocket, core-pivot domain, or dimer interface. Substitutions to the DNA-binding domain or dimer interface generally decrease *EC*_50_. Substitutions to the ligand-binding pocket or core-pivot domain generally increase *EC*_50_.

In addition to specific substitutions that affect both *G*_∞_ and *EC*_50_, we identified nine positions (N125, P127, D149, V192, A194, A245, N246, T276, Q291), where different substitutions either reduce *G*_∞_ by more than 5-fold or increase *EC*_50_ by more than 5-fold, but not both. These positions are all in the ligand-binding pocket. We also identified five positions (H74, V80, K84, S97, M98) where different substitutions reduce either *G*_∞_ or *EC*_50_ by more than 5-fold but not both. These positions are all located at the dimer interface.

Combining multiple substitutions in a single protein almost always has a log-additive effect on *EC*_50_. Only 0.57% (12 of 2101) of double amino acid substitutions have *EC*_50_ values that differ from the log-additive effects of the single substitutions by more than 2.5-fold (Fig. 3). This result, combined with the wide distribution of residues that affect *EC*_50_, suggests that LacI allostery is controlled by a free energy balance with additive contributions from many residues and interactions.

### Phenotypic innovation in an allosteric landscape

Beyond the comprehensive mapping of single-substitution effects, the LacI genotype-phenotype landscape measurement revealed a surprising number of variants with phenotypes that differ qualitatively from the wild type. For example, approximately 230 of the LacI variants have an inverted phenotype (*G*_0_ > *G*_∞_, Fig. 1e), accounting for approximately 0.35% of the measured library (Supplementary Fig. 2a). We verified the dose-response curves for 10 inverted variants with flow cytometry (e.g. Supplementary Fig. 4). To understand the mutational basis for the inverted phenotype, we examined a set of 43 strongly inverted variants (with *G*_0_/*G*_∞_ > 2, *G*_0_ > *G*_∞,*wt*_/2, and *EC*_50_ between 3 µmol/L and 1000 µmol/L). The results indicate that diverse substitutions can lead to the inverted phenotype. For example, we identified 10 amino acid substitutions associated with the inverted phenotype (S70I, K84N, D88Y, V96E, A135T, V192A, G200S, Q248H, Y273H, A343G; Fig. 4a,c). However, none of these substitutions are present in more than 12% of the strongly inverted variants, and 51% of the strongly inverted variants have none of these substitutions. Furthermore, the set of strongly inverted variants are more genetically distant from each other than randomly selected variants from the library (Fig. 4c, Supplementary Fig. 14).

**Fig. 4.**
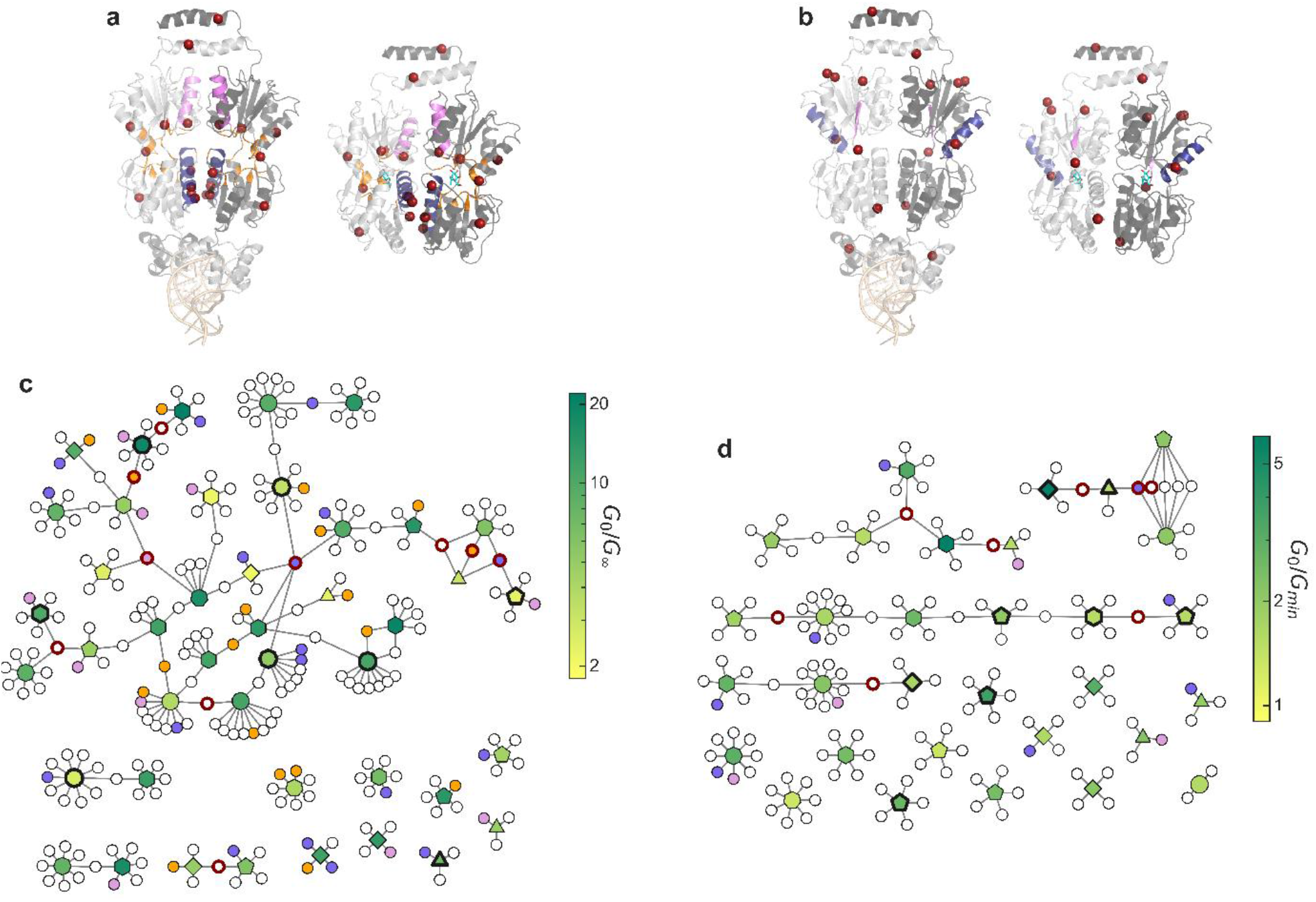
Analysis of inverted and band-stop genotypes. **a-b** Location of amino acid substitutions associated with strongly inverted (**a**) and strong band-stop (**b**) phenotypes. For each plot, the DNA-binding configuration of LacI is shown on the left (PDB ID: 1LGB), with the DNA operator at the bottom in light orange; the ligand-binding configuration is shown on the right (PDB ID: 1LBH), with IPTG in cyan. Both configurations are shown with the view oriented along the protein dimer interface, with one monomer in light gray and the other monomer in dark gray. The locations of associated (i.e. high-frequency) amino acid substitutions are highlighted as red spheres, and secondary structures where inverted or band-stop variants have amino acid substitutions at a significantly higher frequency than the full library are shaded with different colors. For strongly inverted variants (**a**), helix 5 is shaded blue, helix 11 is shaded violet, and the residues near the ligand-binding pocket are shaded orange. For strong band-stop variants (**b**), helix 9 is shaded blue, and strand J is shaded violet. **c-d** Network diagrams showing relatedness among genotypes for strongly inverted (**c**) and strong band-stop (**d**) variants. Within each network diagram, LacI variants are represented by polygonal nodes, with a colormap indicating the *G*_0_/*G*_∞_ or *G*_0_/*G*_*min*_ ratio (see Fig. 1e). The number of sides of the polygon indicates the number of amino acid substitutions relative to the wild-type, and bold outlines indicate variants that were verified with flow cytometry. Smaller circular nodes represent substitutions, with lines showing the substitutions for each variant. Bold red outlines on the substitution nodes indicate the associated substitutions shown as spheres in **a-b**, and the shading of substitution nodes matches the shading used to highlight secondary structures in **a-b**.

The inverted LacI variants can provide specific insight into allosteric biophysics and structure-function relationships, since inversion of the dose-response curve requires inversion of both the allosteric constant^13^ and the relative ligand-binding affinity between the two conformations^2^. Although the set of strongly inverted LacI variants are genetically diverse, many of them have substitutions in similar regions of the protein that may account for the requisite biophysical changes. First, 67% of the strongly inverted variants have substitutions near the ligand-binding pocket (within 7 Å), which likely contribute to the change in ligand-binding affinity. Surprisingly, 21% of the strongly inverted variants have no substitutions within 10 Å of the binding pocket, so binding affinity must be indirectly affected by distal substitutions in those variants. Second, nearly all strongly inverted variants have substitutions at the dimer interface (91%, compared to 54% for the full library), with most (70%) having substitutions in helix 5 (47%), helix 11 (28%), or both (5%, Fig. 4a,c). This suggests that residues in those structural features are important for modulating the allosteric constant.

### Discovery of novel allosteric phenotypes

In addition to the inverted phenotypes, we were surprised to discover LacI variants with dose-response curves that did not match the sigmoidal form of the Hill equation. Specifically, we found variants with band-pass or band-stop dose-response curves, i.e. variants that repress or activate gene expression only over a narrow range of ligand concentrations (e.g. Fig. 1e). Approximately 200 of the LacI variants have band-stop or band-pass phenotypes, accounting for approximately 0.3% of the measured library (Supplementary Fig. 2a). We verified the dose-response curves of 13 band-stop variants and two band-pass variants using flow cytometry (e.g. Supplementary Fig. 5-6). To our knowledge, this is the first identification of single-protein genetic sensors with band-stop dose-response curves.

**Fig. 5.**
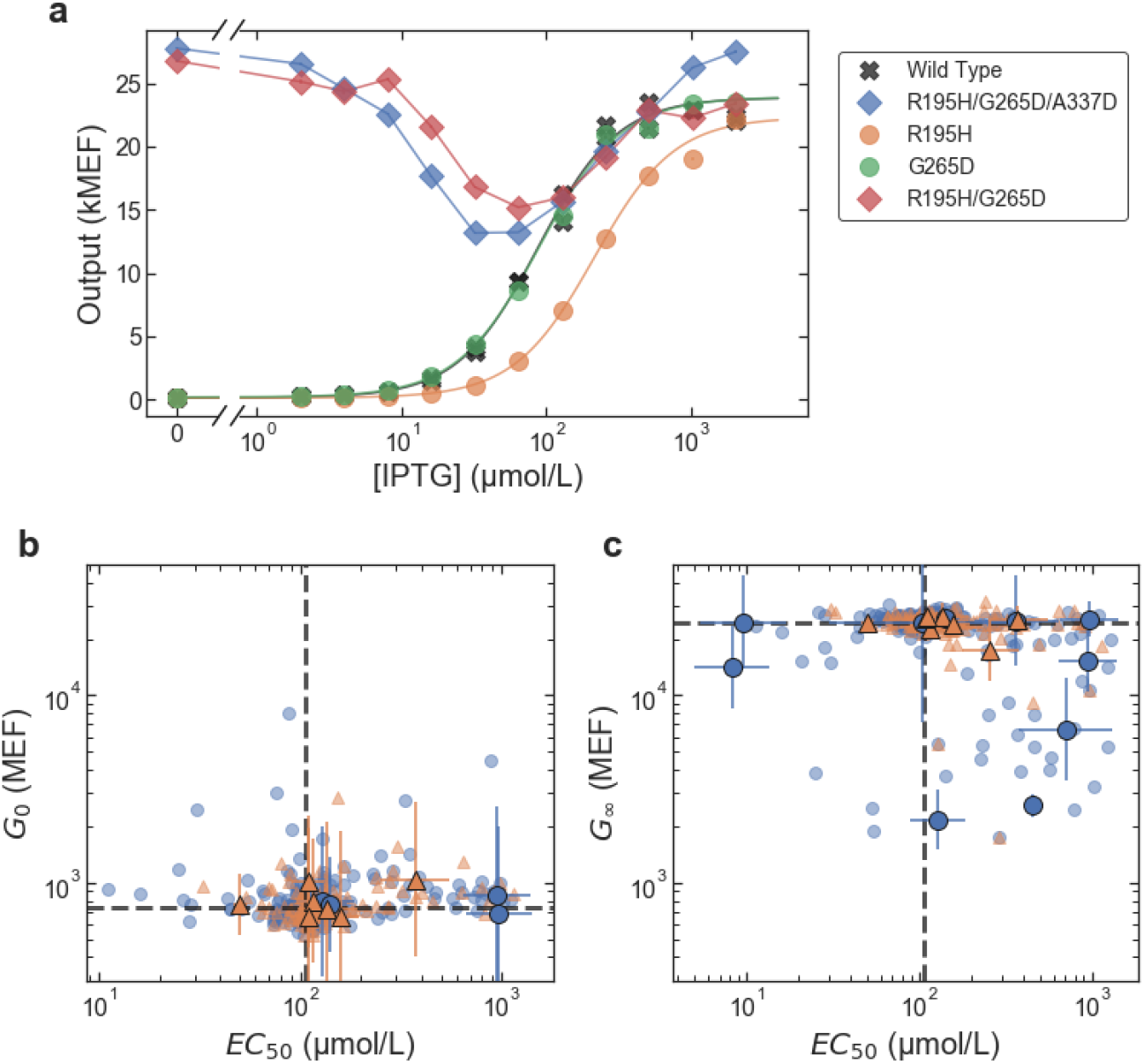
The band-stop phenotype emerges from combinations of nearly silent amino acid substitutions. **a** Dose-response curves measured with flow cytometry for selected LacI variants: wild-type LacI (grey ‘X’s), a strong band-stop variant identified from the library with only three amino acid substitutions (R195H/G265D/A337D; blue diamonds), LacI variants containing the single substitutions R195H (orange circles) and G265D (green circles), LacI variant with the double substitutions R195H/G265D (red diamonds). The single substitution R195H (orange) or G265D (green) results in sigmoidal dose-response curves similar to wild-type LacI, but the combination of the two, R195H/G265D (red), results in a band-stop phenotype. The complete set of permutations of R195H, G265D, and A337D are shown in Supplementary Fig. 15. **b-c** Effects of individual amino acid substitutions associated with inverted and band-stop phenotypes. Each plot shows the joint effect of individual amino acid substitutions on two Hill equation parameters. The blue circles plotted with error bars show the effects of substitutions associated with the strongly inverted phenotype and the orange triangles plotted with error bars show the effects of substitutions associated with the strong band-stop phenotype. Most substitutions associated with the inverted phenotype cause a large shift in either *EC*_50_, *G*_∞_, or both, consistent with the biophysical requirements for inverting the dose-response curve. In contrast, most of the amino acid substitutions associated with the band-stop phenotype are nearly silent. Light blue circles and light orange triangles show the effects for all amino acid substitutions found in the sets of strongly inverted and strong band-stop variants, respectively. Dashed gray lines mark the wild-type parameter values. Plotted data includes a combination of direct experimental measurements and DNN model predictions and is included in Supplementary Data 1. Error bars indicate ± one standard deviation.

Phenotypic similarities between the band-stop and inverted LacI variants (i.e. high *G*_0_, and initially decreasing gene expression as ligand concentration increases) imply similar biophysical requirements. However, amino acid substitutions associated with the band-stop phenotype are remarkably different from those for the inverted phenotype. While inverted variants often have substitutions near the ligand-binding pocket and dimer interface, a set of 31 strong band-stop variants are twice as likely as the full library to have substitutions in helix 9 (32% compared to 16%) and nearly four times as likely to have substitutions in strand J (13% compared to 3.4%). Helix 9 is on the periphery of the protein, and strand J is in the center of the C-terminal core domain. Furthermore, 100% of the strong band-stop variants have substitutions in the C-terminal core of the protein, compared with 79% of the full library (Fig. 4b,d).

To further investigate the band-stop phenotype, we chose a strong band-stop LacI variant with only three amino acid substitutions (R195H/G265D/A337D). We synthesized LacI variants with all possible combinations of those substitutions and measured their dose-response curves with flow cytometry. Although each single substitution resulted in a sigmoidal dose-response similar to wild-type LacI, the combination of two substitutions (R195H/G265D) gave rise to the band-stop phenotype (Fig. 5a, Supplementary Fig. 15). To test whether this result applies to the band-stop phenotype generally, we used the single-substitution effects presented above to examine each of the substitutions associated with the strong band-stop phenotype. Individually, the substitutions associated with the band-stop phenotype are nearly silent, i.e. they have little or no effect on the dose-response curve; yet in combination with other substitutions, they result in the band-stop phenotype. In contrast, most of the individual substitutions associated with the inverted phenotype cause a large shift in either *EC*_50_, *G*_∞_, or both (Fig. 5b,c).

## Discussion

For the goal of an improved understanding of allostery, our results reveal the dual nature of the problem: First, the DNN model and the mapping of single-substitution effects demonstrate that large-scale measurements and analysis can overcome the challenges inherent to the structural complexity of allosteric function. They can provide accurate predictions for specific allosteric proteins and can also reveal systematic structure-function relationships that may be more generalizable (i.e. the importance of the dimer interface and the log-additivity of *EC*_50_). However, the band-stop phenotype highlights the limits of that predictability, as well as the constraints of conventional models of allostery. While the allosteric function of many LacI variants is well-described by the Monod-Wyman-Changeux (MWC) model of allostery^2,13,14^, the band-stop phenotype is inconsistent with that model. In particular, the biphasic dose-response of the band-stop variants suggests negative cooperativity and that the relevant free-energy changes may be more entropic than structural^1^. Our most surprising and unpredictable result is the emergence of the band-stop phenotype from combinations of nearly silent amino acid substitutions. However, with over one hundred genetically diverse band-stop variants, our dataset provides a basis for more systematic understanding even in this case. Furthermore, the relatively high abundance of inverted and band-stop variants (approximately 0.35% and 0.2% of the library, respectively, Supplementary Fig. 2a) with genotypes near the wild-type suggests that allosteric genotype-phenotype landscapes allow for rapid evolutionary innovation, a conclusion that is supported by the existence of natural transcription factors related to LacI with inverted phenotypes^24,25^.

Overall, our findings suggest that a surprising diversity of useful and potentially novel allosteric phenotypes exist with genotypes that are discoverable only via large-scale landscape measurements.

## Methods

### Strain, plasmid, and library construction

All reported measurements were completed using *E. coli* strain MG1655Δ*lac*^26^. Briefly, strain MG1655Δ*lac* was constructed by replacing the lactose operon of *E. coli* strain MG1655 (ATCC #47076) with the bleomycin resistance gene from *Streptoalloteichus hindustanus* (*Shble*).

Two plasmids were used for this work: a library plasmid (pTY1, Supplementary Fig. 1a) used for the measurement of the genotype and phenotype of the entire LacI library, and a verification plasmid (pVER, Supplementary Fig. 1b) used to verify the function of over 100 LacI variants from the library chosen to test the accuracy of the library-scale dose-response curve measurement method. A step-by-step description of the plasmid assembly protocol is available^27^. The sequences are available in GenBank (MT702633, MT702634, for pTY1 and pVER, respectively).

Plasmid pTY1 contained the *lacI* coding DNA sequence (CDS) and the lactose operator (*lacO*) regulating the transcription of a tetracycline resistance gene, *tetA*, which, in the presence of tetracycline, confers a measurable change in fitness connected with the expression level of the regulated genes. Plasmid pTY1 also encoded Enhanced Yellow Fluorescent Protein (YFP), which was used during library construction to select a library in which most of the LacI variants could function as allosteric repressors (see below).

Plasmid pVER contained a similar system in which LacI and *lacO* regulate the transcription of only YFP. Plasmid pVER was used to measure dose-response curves of clonal LacI variants using flow cytometry. Each variant chosen from the library for verification was chemically synthesized (Twist Biosciences), inserted into pVER, and transformed into *E. coli* strain MG1655Δ*lac* for flow cytometry measurements to confirm the dose-response curve inferred from the library-scale measurements.

The LacI library was generated by error-prone PCR of the wild-type *lacI*. The library was inserted into pTY1 along with randomly synthesized DNA barcodes. Each barcode consisted of 54 random nucleotides introduced with PCR primers (Integrated DNA Technologies). Most of the variants in the initial library had high *G*(0), i.e. the *I*^−^ phenotype^18^. To generate a library in which most of the LacI variants could function as allosteric repressors, we used fluorescence activated cell sorting (FACS) to select a portion of the library with low fluorescence in the absence of ligand (Sony SH800S Cell Sorter). To allow comprehensive long-read sequencing of the library (PacBio sequel II, see Long-read sequencing section, below), we further reduced the library size by dilution of the FACS-selected library to create a population bottleneck of the desired size. For the work reported here, we used a library of approximately 10^5^ LacI variants (determined by serial plating and colony counting).

A spike-in control strain was used to normalize the DNA barcode read counts for the sequencing-based fitness measurement (see Library-scale fitness measurement section, below). The spike-in control strain contained the Library Plasmid with a LacI variant that had a constant, high *tetA* expression level. The fitness of the spike-in control was determined from OD_600_ data acquired during growth of clonal cultures with the same automated growth protocol as used for the genotype-phenotype landscape measurement (see Growth protocol for landscape measurement section, below). The fitness of the spike-in control was measured in all 24 chemical environments and was independent of IPTG concentration but was slightly lower with tetracycline (0.75 hour^-1^) than without tetracycline (0.81 hour^-1^).

### Culture conditions

Unless otherwise noted, *E. coli* cultures were grown in a rich M9 media (3 g/L KH2PO4, 6.78 g/L Na2HPO4, 0.5 g/L NaCl, 1 g/L NH4Cl, 0.1 mmol/L CaCl2, 2 mmol/L MgSO4, 4% glycerol, and 20 g/L casamino acids) supplemented with 50 µg/mL kanamycin.

*E. coli* cultures were grown in a laboratory automation system that controlled preparation of 96-well culture plates with media and additives (i.e. IPTG and tetracycline). Cultures were grown in clear-bottom 96-well plates with 1.1 mL square wells (4titude, 4ti-0255). The culture volume per well was 0.5 mL. Before incubation, an automated plate sealer (4titude, a4S) was used to seal each 96-well plate with a gas permeable membrane (4titude, 4ti-0598). Cultures were incubated in a multi-mode plate reader (BioTek, Neo2SM) at 37 °C with a 1 °C gradient applied from the bottom to the top of the incubation chamber to minimize condensation on the inside of the membrane. During incubation, the plate reader was set for double-orbital shaking at 807 cycles per minute. Optical density at 600 nm (OD600) was measured every 5 minutes during incubation, with continuous shaking applied between measurements. After incubation, an automated de-sealer (Brooks, XPeel) was used to remove the gas permeable membrane from each 96-well plate.

### Growth protocol for landscape measurement

To measure the fitness and dose-response curve of every LacI variant in the library, a culture of *E. coli* containing the LacI library was mixed at a 99:1 ratio with a culture of the *E. coli* spike-in control. The culture was loaded into the automated microbial growth and measurement system where it was distributed across a 96-well plate and then grown to stationary phase (12 hours). Cultures were then diluted 50-fold into a new 96-well plate, Growth Plate 1, containing 11 rows with a 2-fold serial dilution gradient of IPTG with concentrations ranging from 2 µmol/L to 2048 µmol/L and one row without IPTG. Growth in IPTG allowed each variant to reach a steady state tetA expression level in each IPTG concentration. Growth Plate 1 was grown for 160 minutes, corresponding to approximately 3.3 generations, and then diluted 10-fold into Growth Plate 2. Growth Plate 2 contained the same IPTG gradient as Growth Plate 1 with the addition of tetracycline (20 µg/mL) to alternating rows in the plate, resulting in 24 chemical environments, with 4 duplicate wells for each environment. Growth Plate 2 was grown for 160 minutes and then diluted 10-fold into Growth Plate 3, which contained the same 24 chemical environments as Growth Plate 2. This process was repeated for Growth Plate 4, which also contained the same 24 chemical environments. The total growth time for the fitness measurements in the 24 chemical environments, 480 minutes across Growth Plates 2-4, corresponded to approximately 10 generations for the fastest-growing cultures. The 50-fold dilution factor from stationary phase into Growth Plate 1 and the 160 minute growth time per plate were chosen to maintain the cultures in exponential growth for the entire 480 minutes. During each 160 minute incubation, the cultures without tetracycline increased approximately 10-fold in optical density, to a final OD_600_ of approximately 0.5 (corresponding to an estimated cell density of 4 × 10^8^ cells/mL).

After each growth plate was used to seed the subsequent plate (or at the end of 160 minutes for Growth Plate 4), the remaining culture volumes for each chemical environment (approximately 450 µL/well, four duplicates per plate) were combined and pelleted by centrifugation (3878 *g* for 10 minutes at 23 °C). Plasmid DNA was then extracted from the 24 combined samples with a custom method using reagents from the QIAprep Miniprep Kit (Qiagen cat. #27104) on an automated liquid handler equipped with a positive-pressure filter press (step-by-step protocol available^28^). After extraction, DNA was eluted into a final volume of approximately 50 µL and the concentration of DNA in each sample ranged from undetectable up to approximately 1.5 ng/µL. This corresponds to an estimated maximum of approximately 10^10^ plasmids per sample.

### Barcode sequencing

After plasmid extraction, each set of 24 plasmid DNA samples was prepared for barcode sequencing using a custom sequencing sample preparation method on a second automated liquid handler (step-by-step protocol is available^29^). Briefly, the plasmid DNA was linearized with ApaI restriction enzyme. Then, a 3-cycle PCR was performed to attach sample multiplexing tags to the resulting amplicons so the different samples could be distinguished when pooled and run on the same sequencing flow cell. Eight forward index primers and 12 reverse index primers were used to label the amplicons from each sample across the 24 chemical environments and the four time points. After a magnetic-bead-based cleanup step, a second, 15-cylce PCR was run to attach the standard Illumina paired-end adapter sequences and to amplify the resulting amplicons for sequencing. After a second magnetic-bead-based cleanup, the 24 samples from each time point were pooled and stored at 4 °C until sequencing. For sequencing, DNA was diluted to a final concentration of approximately 5 nmol/L and combined with 20% phiX control DNA. DNA from each of the 4 time points was sequenced in a separate lane on an Illumina HiSeqX using paired-end mode with 150 bp in each direction.

To count DNA barcodes and estimate the fitness associated with each LacI variant, the sequencing data was analyzed using custom software written in C# and Python, and the Bartender1.1 barcode clustering algorithm^30^ (https://github.com/djross22/nist_lacI_landscape_analysis).

The sequence of the nominal Illumina compatible amplicon was (with Illumina adapters and flow cell binding sequences in gray):

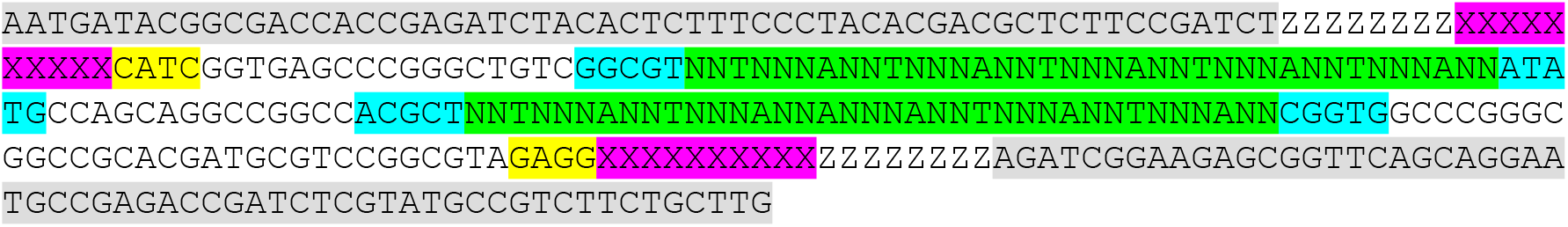

The nominal forward and reverse reads from paired-end barcode sequencing were:

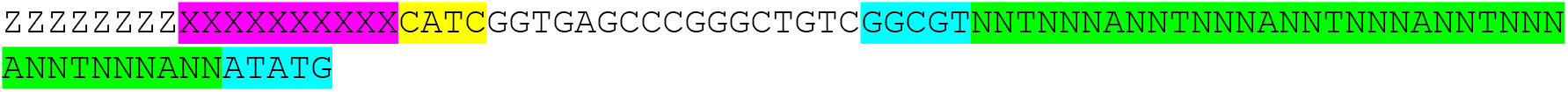

and

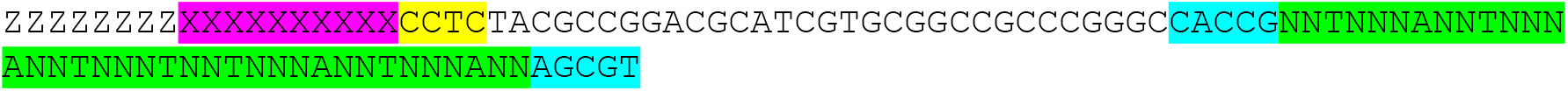

The Z’s at the beginning of each read are random nucleotides used as unique molecular identifiers (UMIs) to correct for PCR jackpotting^31^, the X’s are the sample multiplexing tag sequences, and the N’s are the random nucleotides of the DNA barcodes. To minimize the chances of barcode crosstalk, we used dual barcodes, with independent random barcode sequences on the forward and reverse reads and 27 random nucleotides in each of the forward and reverse barcodes.

The raw sequences were parsed, and sequences were kept for further analysis only if they passed the following quality criteria for both the forward and reverse reads:

1. The four bases after the multiplexing tag (highlighted in yellow above) must match the nominal sequence with one allowed mismatch, and the multiplexing tag sequence (highlighted in pink above) must match the nominal sequence for one of the multiplexing tags used with up to three allowed mismatches.
2. The five flanking bases before and after the barcodes (highlighted in cyan above) must match the nominal sequence with one allowed mismatch per set of five bases, and the number of bases in the barcode (highlighted in green above) must be between 35 and 41 (inclusive).
3. The mean Illumina quality score for the barcode and the five flanking bases before and after the barcode must be greater than 30.

For the four lanes of HiSeq data, there were 2,024,537,456 raw reads, of which 1,576,168,836 reads passed the quality criteria (78%). Note that 20% of the DNA sample loaded onto the HiSeq instrument was phiX DNA.

True barcode sequences were identified using the Bartender1.1 clustering algorithm^30^ with the following parameter settings: maximum cluster distance = 4, cluster merging threshold = 8, cluster seed length = 5, cluster seed step = 1, frequency cutoff = 500. Barcodes from the forward and reverse reads were clustered independently. The Bartender1.1 clustering algorithm identified 43,259 distinguishable forward barcode clusters and 31,055 distinguishable reverse barcode clusters.

To correct for insertion-deletion read errors, barcode clusters of different length were considered for merging. First, barcode clusters with sequences that were sub-strings of one another were automatically merged. Second, pairs of barcode clusters with a DNA sequence Levenshtein distance of 1 or 2 were merged if the ratio of the smaller cluster read count to the total read count of both clusters was less than 0.001 and 0.0001, respectively. Third, all barcode clusters with a Levenshtein distance less than 7 from the barcode for the spike-in control were merged.

After merging barcode clusters of different lengths, there were 43,169 distinguishable forward barcode clusters and 30,931 distinguishable reverse barcode clusters. The random positions within the forward and reverse barcodes had approximately equal probabilities for each nucleotide, with a mean entropy per position of 1.9799 bits ± 0.0066 bits.

After barcode clustering and merging, the barcode sequencing reads were sorted based on the sample multiplexing tags and the barcode read counts were corrected for PCR jackpotting effects. Sets of multiple barcode reads were treated as PCR jackpot duplicates if they had the same UMI sequence, the same multiplexing tag, and the same barcode sequence for both forward and reverse barcode reads. In the corrected barcode count, each set of PCR jackpot duplicates was counted as a single read. Approximately 15% of the total barcode sequencing reads were found to be PCR jackpot duplicates.

The forward and reverse barcodes were then combined to give the DNA barcodes used to measure the relative abundance of each LacI variant in the library. An additional barcode count threshold was applied, keeping only DNA barcodes with a total read count (across all 24 environments and 4 time points) greater than 2000. A small number (139) of DNA barcodes were identified as likely chimeras with forward and reverse barcodes combined from different plasmid templates^32–34^. The likely chimera barcodes were not used in further analysis.

Finally, 14 pairs of DNA barcodes were found with DNA sequence Hamming distance of one (across both forward and reverse barcodes). Only one DNA barcode from each pair was also found in the long-read sequencing data (see Long-read sequencing section, below). In addition, the fitness curves (vs. IPTG concentration) were very similar for both barcodes in each pair. Based on this, the read counts associated with each of those 14 pairs of dual barcodes were merged, and each pair was treated as a single DNA barcode.

The final set of 67,730 DNA barcodes was used for all subsequent analysis to extract estimates of the fitness and dose-response curve associated with each barcode.

### Long-read sequencing

The full sequence of the Library Plasmid for every LacI variant in the library was measured using PacBio circular consensus HiFi sequencing. The HiFi sequencing data was used to determine the consensus *lacI* sequence for each variant and the corresponding DNA barcode. Of the 67,731 distinct DNA barcodes (see Barcode sequencing section, above), the HiFi sequencing data was used to determine the *lacI* sequences for 63,064 (93%), 3,878 with a single HiFi lacI read, and 59,186 with multiple HiFi lacI reads.

In addition, the full plasmid sequence was used to detect unintended mutations in the plasmid, i.e. mutations to plasmid regions other than the *lacI* CDS. For analysis of the HiFi read data, the full plasmid sequence was divided into 11 non-overlapping regions that roughly correspond to different functional elements of the plasmid (Supplementary Table 1), and the sequences for each region were extracted from the HiFi reads using a custom bioinformatic pipeline (https://github.com/djross22/nist_lacI_landscape_analysis). The number of unintended mutations to plasmid regions other than the *lacI* CDS was relatively low (Supplementary Table 1), so it was not possible to examine mutational effects with base-pair- or residue-level resolution. However, by pooling the mutational information for each region, significant region-specific effects could be detected. To determine if mutations in a region of the plasmid had a significant effect, the estimated Hill equation parameters were compared for all variants with one or more mutations in a given plasmid region vs. all variants with zero mutations in that region. Significant differences in the geometric mean of one or more Hill equation parameters were found for variants with mutations in the following regions: tetA (p-value for log_10_(*G*_∞_): 2 × 10^−56^), KAN (p-value for log_10_(*G*_∞_): 4 × 10^−11^), origin of replication (p-value for log_10_(*G*_∞_): 6 × 10^−14^), and YFP (p-value for log_10_(*G*_0_): 4 × 10^−109^; p-value for log_10_(*G*_∞_): 5 × 10^−10^; p-value for log_10_(*EC*_50_): 2 × 10^−74^), where the p-values given are for Welch’s unequal-variances t-test.

In addition, 43 of the 535 variants with the wild-type LacI amino acid sequence had mutations in the regulatory region (containing the P_lacI_ and P_tacI_ promoters, the *lacO* operator, the riboJ insulator, and the RBS sites for both *lacI* and *tetA*). Of those 43 variants, 3 had *EC*_50_ values that differed by approximately 2-fold or more from the geometric mean value for the wild-type *EC*_50_. The Kolmogorov-Smirnov test was used to compare the distributions of *EC*_50_ values between the wild-type variants with and without mutations in the regulatory region; the results indicated a significant difference (p-value: 0.024).

To avoid biasing the results of the machine learning and other quantitative phenotypic analyses, variants were excluded from those analyses if they had one or more mutations in the non-*lacI* regions that show significant mutational effects: tetA, YFP, KAN, the origin of replication, and the regulatory region. After applying this data quality filter in addition to those described above, there were 54,162 variants that we used for further quantitative analysis.

### Library-scale fitness measurement

The experimental approach for this work was designed to maintain bacterial cultures in exponential growth phase for the full duration of the measurements. So, in all analysis, the Malthusian definition of fitness was used, i.e. fitness is the exponential growth rate^35^.

The fitness of cells containing each LacI variant was calculated from the change in the relative abundance of DNA barcodes over time. The spike-in control was used to normalize the DNA barcode count data to enable the determination of the absolute fitness for each LacI variant in the library.

Briefly, for each LacI variant in each of the 24 chemical environments, the ratio of the barcode read count to the spike-in read count was fit to a function assuming exponential growth and a lag in the onset of the fitness impact of tetracycline. The fitness associated with each variant in each of the 24 chemical environments was determined as a parameter in the corresponding least-squares fit as detailed below.

The barcode sequencing data was analyzed with a model based on the assumption that the number of cells containing each LacI variant grows with an exponential expansion rate that is independent of all other variants. So, for each sample, at the end of the incubation cycle for Growth Plate *j*, the number of cells with LacI variant *i* is:

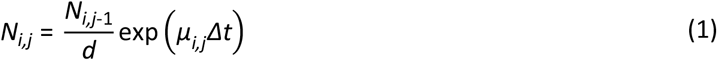

where, *d* (= 10) is the dilution factor used in transferring the cell culture from Growth Plate *j* – 1 to Growth Plate *j, Δt* (≈ 165 minutes) is the total incubation time for each growth plate (including time required for automated cell passaging), and *μ*_*i,j*_ is the fitness (ie mean exponential growth rate) of cells with LacI variant *i* in Growth Plate *j*.

For samples without tetracycline, the chemical composition of the media was the same for all growth plates, so the fitness is assumed to be constant, *μ*_*i,j*_ = *μ*_*i*_^0^, where *μ*_*i*_^0^ is the fitness associated with LacI variant *i* in the absence of tetracycline. Consequently, the number of cells in each Growth Plate for samples grown without tetracycline is:

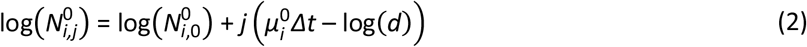

where 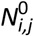 is the number of cells with LacI variant *i* at the end of Growth Plate *j* for samples grown without tetracycline.

For samples grown with tetracycline, the tetracycline was only added to the culture media for Growth Plates 2-4. Because of the mode of action of tetracycline (inhibition of translation), there was a lag in its effect on cell fitness. Accordingly, the analysis assumes fitness varies as a function of time:

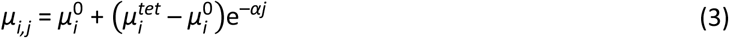

where *μ*_*i*_^*tet*^ is the steady-state fitness with tetracycline, and *α* is a transition rate. Based on test measurements with a small-scale library, the transition rate was kept fixed at *α* = log(5). From Eq. (3), the number of cells in each Growth Plate for samples grown with tetracycline is:

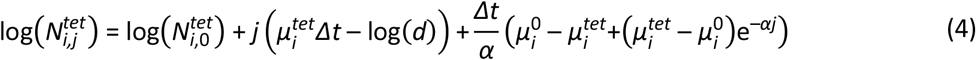

The barcode read count for variant *i* in Growth Plate *j* was assumed to be proportional to the cell number:

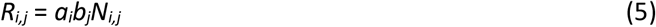

where *a*_*i*_ is a proportionality constant associated with variant *i*, and *b*_*j*_ is a proportionality constant associated with Growth Plate *j*. The proportionality constant *a*_*i*_ can be different for each variant *i* due to differences in PCR amplification efficiency resulting from variations in the barcode sequences on each amplicon. Similarly, the proportionality constant *b*_*j*_ can be different for each Growth Plate because of sample-to-sample variations in the DNA extraction efficiency or differences in PCR efficiency associated with different sample multiplexing tag sequences.

The logarithm of the read count normalized by the spike-in read count was used to estimate the fitness of each variant from its associated barcode read count:

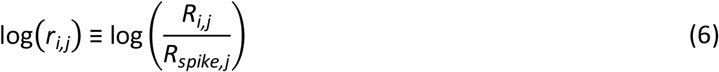

For samples without tetracycline, *μ*_*i*_^0^ was estimated for each variant using a weighted linear least-squares fit to the log-count ratio vs. *j*:

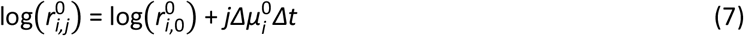

where 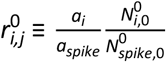, and 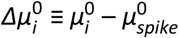 is the difference between the fitness of variant *i* and the spike-in fitness without tetracycline.

For samples grown with tetracycline, *μ*_*i*_^*tet*^ was estimated for each variant with a weighted least-squares fit to the non-linear form for the log-count ratio:

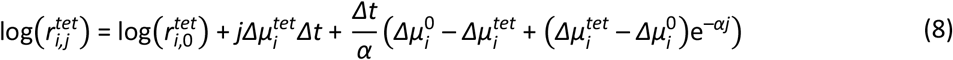

where 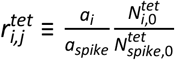, and 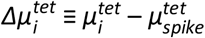 is the difference between the fitness of variant *i* and the spike-in fitness with tetracycline.

For the least-squares fits to determine both *μ*_*i*_^0^ and *μ*_*i*_^*tet*^, the fits were weighted based on the propagated uncertainties of 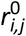 and 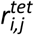 calculated assuming that the uncertainty of each read count was dominated by Poisson sampling.

For the fitness landscape measurement, there were a large number of outliers for the read count measurements from three of the samples: Growth Plate 3, without tetracycline, [IPTG] = 8 µmol/L; Growth Plate 4, without tetracycline, [IPTG] = 64 µmol/L and [IPTG] = 2048 µmol/L. These three samples were excluded from the analysis.

### Dose-response curve measurements

Plasmids pTY1 and pVER were engineered to provide two independent measurements of the dose-response curve for LacI variants. First, in pTY1, LacI regulates the expression of a tetracycline resistance gene (*tetA*) that enables determination of the dose-response from barcode sequencing data by comparing the fitness measured with tetracycline to the fitness measured without tetracycline. Second, in pVER, the LacI regulates the expression of a fluorescent protein (YFP) that enables direct measurement of the dose-response curve with flow cytometry.

A set of nine randomly selected LacI variants were used to calibrate the estimation of regulated gene expression output from the barcode-sequencing fitness measurements (Supplementary Fig. 16). The calibration data consisted of the fitness data for each calibration variant from the library barcode sequencing measurement (using the library plasmid, pTY1) and flow cytometry data for each calibration variant prepared as a clonal culture (using the verification plasmid, pVER). This data was fit to a Hill equation model for the fitness impact of tetracycline as a function of the regulated gene expression level, *G*:

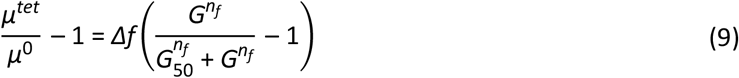

where *µ*^*tet*^ is the fitness with tetracycline, *µ*^0^ is the fitness without tetracycline, *Δf* is the maximal fitness impact of tetracycline (when *G* = 0), *G*_50_ is the gene expression level that produces a 50% recovery in fitness, and *n*_*f*_ characterizes the steepness of the fitness calibration curve. Because the fitness calibration curve, Eq. (9), is nonlinear, it cannot be directly inverted to give the regulated gene expression level for all possible fitness measurements. So, two Bayesian inference models were used to estimate the dose-response curves for every LacI variant in the library using the barcode sequencing fitness measurements. Source code for both models is included in the software archive at https://github.com/djross22/nist_lacI_landscape_analysis. Both inference models used Eq. (9) to represent the relationship between fitness and regulated gene expression. The parameters *Δf, G*_50_, and *n*_*f*_ were included in both inference models as parameters with informative priors. Priors for *G*_50_ and *n*_*f*_ were based on the results of the fit to the fitness calibration data (Supplementary Fig. 16: *G*_50_ ∼ normal(mean=13,330, std=500), *n*_*f*_ ∼ normal(mean=3.24, std=0.29). We chose the prior for *Δf* based on an examination of *μ*^*tet*^/*μ*^0^ − 1 measured with zero IPTG: *Δf* ∼ exponentially-modified-normal(mean=0.720, std=0.015, rate=14). The use of a prior for *Δf* with a broad right-side tail was important to accommodate variants in the library for which *μ*^*tet*^/*μ*^0^ − 1 was systematically less than -0.722.

The first Bayesian inference model assumed that the dose-response curve for each LacI variant was described by the Hill equation. The Hill equation parameters for each variant, *G*_∞_, *G*_0_, *EC*_50_, and *n* and their associated uncertainties were determined using Bayesian parameter estimation by Markov Chain Monte Carlo (MCMC) sampling with PyStan^36^. Broad, flat priors were used for log_10_(*G*_0_), log_10_(*G*_∞_), and log_10_(*EC*_50_), with error function boundaries to constrain those parameter estimates to within the measurable range (100 MEF ≤ *G*_0_, *G*_∞_ ≤ 50,000 MEF; 0.1 µmol/L ≤ *EC*_50,*i*_ ≤ 40,000 µmol/L). The prior for *n*_*i*_ was a gamma distribution with shape parameter of 4.0 and inverse scale parameter of 3.33. The inference model was run individually for each LacI variant, with four independent chains, 1000 iterations per chain (500 warmup iterations), and the adapt_delta parameter set to 0.9. Testing with data from a set of randomly selected variants indicated that these settings for the Stan sampling algorithm typically produced a Gelman-Rubin 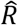 diagnostic less than 1.05 and number of effective iterations greater than 100.

The second Bayesian inference model was a non-parametric Gaussian process (GP) model^37^ that assumed only that the dose-response curve for each LacI variant was a smooth function of IPTG concentration. The GP model was used to determine which variants had band-pass or band-stop phenotypes. The GP model was also implemented using MCMC sampling with PyStan^36^. The GP inference model was run individually for each variant, with four independent chains, 1000 iterations per chain (500 warmup iterations), and the adapt_delta parameter set to 0.9. Testing with data from a set of randomly selected variants indicated that these settings for the Stan sampling algorithm of the GP model typically produced a Gelman-Rubin 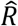 diagnostic less than 1.02 and number of effective iterations greater than 200.

### Flow cytometry measurements

Over 100 LacI variants from the library were chosen for flow cytometry verification of the dose-response curves. The CDSs of these variants were chemically synthesized (Twist Bioscience), cloned into the verification plasmid, pVER, and then transformed into MG1655Δ*lac*. Transformants were plated in LB supplemented with kanamycin and 0.2% glucose. LacI variant sequences were verified with Sanger sequencing (Psomagen USA). For flow cytometry measurements of dose-response curves, a culture of *E. coli* containing pVER with a chosen variant sequence was distributed across 12 wells of a 96-well plate and grown to stationary phase using the automated microbial growth system. After growth to stationary phase, cultures were diluted 50-fold into a plate containing the same 12 IPTG concentrations used during the fitness landscape measurement (0 µmol/L to 2048 µmol/L). In some cases, higher IPTG concentrations were used to capture the full dose-response curves of selected variants (e.g. Supplementary Figs. 4-5). Cultures were then grown for 160 minutes (∼3.3 generations) before being diluted 10-fold into the same IPTG gradient and grown for another 160 minutes. Then, 5 µL of each culture was diluted into 195 µL of PBS supplemented with 170 µg/mL chloramphenicol and incubated at room temperature for 30-60 minutes to halt the translation of YFP and allow extant YFP to mature in the cells.

Samples were measured on an Attune NxT flow cytometry with autosampler using a 488 nm excitation laser and a 530 nm ± 15 nm bandpass emission filter. Blank samples were measured with each batch of cell measurements, and an automated gating algorithm was used to discriminate cell events from non-cell events (Supplementary Fig. 17a-b). With the Attune cytometer, the area and height parameters for each detection channel are calibrated to give the same value for singlet events. So, to identify singlet cell events and exclude multiplet cell events, a second automated gating algorithm was applied to select only cells with side scatter area ≅ side scatter height (Supplementary Fig. 17c-d). All subsequent analysis was performed using the singlet cell event data. Fluorescence data was calibrated to molecules of equivalent fluorophore (MEF) using fluorescent calibration beads (Spherotech, part no. RCP-30-20A). The cytometer was programmed to measure a 25 µL portion of each cell sample, and the 40-fold dilution used in the cytometry sample preparation resulted in approximately 20,000 singlet cell measurements per sample. The geometric mean of the YFP fluorescence was used as a summary statistic to represent the regulated gene expression level as a function of the input ligand concentration, [IPTG] for each LacI variant.

### Calculation of abundance for LacI phenotypes

The relative abundance of the various LacI phenotypes (Supplementary Fig. 2) was estimated using the results of both Bayesian inference models (Hill equation and GP). Variants were labeled as “flat response” if the Hill equation model and the GP model agreed (i.e. if the median estimate for the Hill equation dose-response curve was within the central 90% credible interval from the GP model at all 12 IPTG concentrations) and if the posterior probability for *G*_0_ > *G*_∞_ was between 0.05 and 0.95 (from the Hill equation model inference). Variants were labeled as having a negative response if the slope, ∂*G*/∂*L*, was negative at one or more IPTG concentrations with 0.95 or higher posterior probability (from the GP model inference). To avoid false positives from end effects, this negative slope criteria was only applied for IPTG concentrations between 2 µmol/L IPTG and 1024 µmol/L. Variants were labeled as “always on” (the *I*^*–*^ phenotype from reference^18^) if they were flat-response and if *G*(0) was greater than 0.25 times the wild-type *G*_∞_ value with 0.95 or higher posterior probability (from the GP model inference). Variants were labeled as “always off” (the *I*^*S*^ phenotype from reference^18^) if they were flat-response but not always on. Variants were labeled as band-stop or band-pass if the slope, ∂*G*/∂*L*, was negative at some IPTG concentrations and positive at other IPTG concentrations, both with 0.95 or higher posterior probability (from the GP model inference). Band-stop and band-pass variants were distinguished by the ordering of the negative-slope and positive-slope portions of the dose-response curves. Variants that had a negative response but that were not band-pass or band-stop, were labeled as inverted. False-positive rates were estimated for each phenotypic category by manually examining the fitness vs. IPTG data for LacI variants with less than three substitutions. Typical causes of false-positive phenotypic labeling included unusually high noise in the fitness measurement and biased fit results due to outlier fitness data points. Estimated false-positive rates ranged between 0.001 and 0.005. The relative abundance values shown in Supplementary Fig. 2a were corrected for false positives using the estimated rates.

### Comparison of synonymous mutations

The library contained a set of 39 variants with the wild-type *lacI* CDS (but different DNA barcodes), and a set of 310 variants with only synonymous nucleotide changes (*i*.*e*. no amino acid substitutions). Both sets had long-read sequencing coverage for the entire plasmid and were screened to retain only variants with zero unintended mutations in the plasmid (i.e. no mutations in regions of the plasmid other than the *lacI* CDS). The Hill equation fit results for those two sets were compared to determine whether synonymous nucleotide changes significantly affected the phenotype. The Kolmogorov-Smirnov test was used to compare the distributions of Hill equation parameters between these two sets. The resulting p-values (0.71, 0.40, 0.28, and 0.17 for *G*_0_, *G*_∞_, *EC*_50_, and *n* respectively) indicate that there were no significant differences between them. Additionally, the library contained 40 sets of variants, each with four or more synonymous CDSs (including the set of synonymous wild-type sequences and 39 non-wild-type sequences). A hierarchical model was used to compare the Hill equation parameters within each set of synonymous CDSs. Within each set, the uncertainty associated with individual variants was typically larger than the variant-to-variant variability estimated by the hierarchical model. Overall, these results indicate that synonymous SNPs did not measurably impact the LacI phenotype, so only the amino acid sequences were considered for any subsequent quantitative genotype-to-phenotype analysis.

### Analysis of single-substitution data

The single amino acid substitution results presented in Fig. 2, Fig. 5b-c, Supplementary Fig. 13, and included in Supplementary Data 1 are a combination of direct experimental observations, DNN model results, and estimates of *G*_0_ for missing substitutions.

For direct experimental observations, multiple LacI variants were often present in the library with the same single substitution. To ensure that the highest quality data was used for the single-substitution analysis, only data for variants with more than 5000 total barcode reads were used (see Barcode sequencing section, above). For each single substitution, if there was only one LacI variant with more than 5000 barcode reads, the median and standard deviation for each parameter were used directly from the Bayesian inference using the Hill equation model. If there was more than one LacI variant with a given single substitution and more than 5000 barcode reads, the consensus Hill equation parameter values and standard deviations for that substitution were calculated using a hierarchical model based on the eight schools model^38,39^. The hierarchical model was applied separately for each Hill equation parameter. The logarithm of the parameter values was used as input to the hierarchical model, and the input data were centered and normalized by 1.15 × the minimum measurement uncertainty. The standard normal distribution was used as a loosely informative prior for the consensus mean effect, and a half-normal prior (mean = 0.5, std = 1) was used for the normalized consensus standard deviation (i.e. hierarchical standard deviation). These priors and normalization were chosen so that the model gave intuitively reasonable results for the consensus of two LacI variants (i.e. close to the results for the LacI variant with the lowest measurement uncertainty). Results for the hierarchical model were determined using Bayesian parameter estimation by Markov Chain Monte Carlo (MCMC) sampling with PyStan^36^. MCMC sampling was run with 4 independent chains, 10,000 iterations per chain (5,000 warmup iterations), and the adapt_delta parameter set to 0.975.

For *G*_0_, the direct experimental results were used for the 1047 substitutions plotted as gray points or red points and error bars in Fig. 2d and Supplementary Fig. 13. In addition, estimated values were used for the 83 missing substitutions that have been previously shown to result in an “always on” LacI phenotype (i.e., the *I*^*–*^ phenotype ^18,19^). For these substitutions, plotted as pink-gray points and error bars in Fig. 2d, the median value was estimated to be equal to the wild-type value for *G*_∞_ (24,000 MEF), and the geometric standard deviation was estimated to be 4-fold, both based on information from previous publications^18,19^. Note that these 83 substitutions are completely missing from the experimental landscape dataset, i.e. they are not found in any LacI variant, as single substitutions or in combination with other substitutions.

For *G*_∞_ and *EC*_50_, the direct experimental results were used for the 964 substitutions that are found as single substitutions in the library and that have a consensus standard deviation for log_10_(*EC*_50_) less than 0.35. An additional 74 substitutions are found as single substitutions in the library, but with higher *EC*_50_ uncertainty. For these substitutions, either *EC*_50_ is comparable to or higher than the maximum ligand concentration used for the measurement (2048 µmol/L IPTG), or *G*_∞_ is comparable to *G*_0_ (or both). Consequently, the dose-response curve is flat or nearly flat across the range of concentrations used, and the Bayesian inference used to estimate the Hill equation parameters results in *EC*_50_ and *G*_∞_ estimates with large uncertainties. The DNN model can provide a better parameter estimate for these flat-response variants because it uses data and relationships from the full library (e.g. the log-additivity of *EC*_50_) to predict parameter values for each single substitution. So, the DNN model results were used for these 74 substitutions. Finally, the DNN model results were used for an additional 953 substitutions that are found in the library, but only in combination with other substitutions (i.e. not as single substitutions).

### Identification of high-frequency substitutions and structural features associated with inverted and band-stop phenotypes

The set of 43 strongly inverted LacI variants discussed in the main text and used for the plots in Fig. 4a,c were identified by the following criteria: *G*_0_/*G*_∞_ ≥ 2, *G*_0_ > *G*_∞,*wt*_/2, *G*_∞_ < *G*_∞,*wt*_/2, and *EC*_50_ between 3 µmol/L and 1000 µmol/L. The set of 31 strong band-stop variants discussed in the main text and used for the plots in Fig. 4b,d were identified by the following criteria: *G*_0_ > *G*_∞,*wt*_/2, *G*_*min*_ < *G*_∞,*wt*_/2, and the slope, ∂log(*G*)/∂log(*L*), of less than -0.07 at low IPTG concentrations and greater than zero at higher IPTG concentrations, both with 0.95 or higher posterior probability (from the GP model inference). In addition, the sets of strongly inverted and strong band-stop variants were manually screened for likely false positives due to outlier fitness data points.

A hypergeometric test was used to determine the amino acid substitutions that occur more frequently in the set of strongly inverted or strong band-stop variants than in the full library. For each possible substitution, the cumulative hypergeometric distribution was used to calculate the probability of the observed number of occurrences of that substitution in the set of inverted or band-stop variants under a null model of no association. This probability was used as a p-value for the null hypothesis that the observed number of inverted or band-stop variants with that substitution resulted from an unbiased random selection of variants from the full library. Substitutions were considered to occur at significantly higher frequency if they had a p-value less than 0.005 and if they occurred more than once in the set of inverted or band-stop variants. In the set of strongly inverted variants, ten associated (higher frequency) amino acid substitutions were identified: S70I, K84N, D88Y, V96E, A135T, V192A, G200S, Q248H, Y273H, and A343G. In the set of strong band-stop variants, eight associated substitutions were identified: V4A, A92V, H179Q, R195H, G178D, G265D, D292G, and R351G. To estimate the number of false-positives, random sets of LacI variants were chosen with the same sample size as the strongly inverted (43) or the strong band-stop (31) variants and the same significance criteria was applied. From 300 independent iterations of the random selection, the estimated mean number of false-positive substitutions was 2.1 and 2.3 for the inverted and band-stop phenotypes, respectively.

A similar procedure was used to determine which structural features within the protein are mutated with higher frequency in the inverted or band-stop LacI variants. The structural features considered were the secondary structures from the complete crystal structure of LacI^22^, as well as larger structural features (N-terminal core domain, C-terminal core domain, DNA-binding domain, dimer interface) and functional domains (ligand-binding, core-pivot). The p-value threshold used for significance was 0.025. For the strongly inverted variants, six domains were identified with a higher frequency of amino acid substitutions: the dimer interface, residues within 7 Å of the ligand-binding pocket, helix 5, helix 11, strand I, and the N-terminal core. For the strong band-stop variants, three features were identified: the C-terminal core, strand J, and helix 9. From 300 independent random selections of variants from the full library, the estimated mean number of false-positive features was 0.39 and 0.50 for the inverted and band-stop phenotypes, respectively.

### Deep neural network (DNN) modeling

The dataset was pruned to a set of high-quality sequences for DNN modeling. Specifically, data for a LacI variant was only used for modeling if it satisfied the following criteria:

1. No mutations were found in the long-read sequencing results for the regions of the plasmid encoding kanamycin resistance, the origin of replication, the tetA and YFP genes, and the regulatory region containing the promoters and ribosomal binding sites for *lacI* and tetA (Supplementary Table 1).
2. The total number of barcode read counts for a LacI variant was greater than 3000.
3. The number of amino acid substitutions was less than 14.
4. The measurement uncertainty for log_10_(*G*_∞_) was less than 0.7.
5. The results of the Hill equation model and the GP model agreed at all 12 IPTG concentrations. More specifically, data were only used if the median estimate for the dose-response curve from the Hill equation model was within the central 90% credible interval from the GP model at all 12 IPTG concentrations.

After applying the quality criteria listed above, 47,462 LacI variants remained for DNN modeling. The data were used to train the DNN model to predict the Hill equation parameters *G*_0_, *G*_∞_, and *EC*_50_ as detailed below.

Amino acid sequences were represented as one-hot encoded vectors of length L = 2536, and with mutational paths represented as K × L tensors for a sequence with K substitutions. The logarithm of the Hill equation parameter values were normalized to a standard deviation of 1, and then shifted by the corresponding value of the wild-type sequence in order to correctly represent the prediction goal of the change in each parameter relative to wild-type LacI. A long-term, short-term recurrent neural network was selected for the underlying model^17^, with 16 hidden units, a single hidden layer, and hyperbolic tangent (tanh) non-linearities. Inference was performed in pytorch^40^ using the Adam optimizer^41^. For *EC*_50_ and *G*_0_, the contribution of individual data points to the regression loss were weighted inversely proportional to their experimental uncertainty. Model selection was performed with 10-fold cross-validation on the training set (80% of all available data). Approximate Bayesian inference was performed with the Bayes-by-backprop approach^42^. Briefly, this substitutes the point-estimate parameters of the neural network with variational approximations to a Bayesian model, represented as a mean and variance of a normal random variable. Effectively, this only doubles the number of parameters in the model. A mixture of two normal distributions was used as a prior for each parameter weight, with the two mixture components having high and low variance respectively. This prior emulates a sparsifying spike-slab prior while remaining tractable for inference based on back-propagation. Posterior means of each weight were used to calculate posterior predictive means, while Monte-Carlo draws from the variational posterior were used to calculate the model prediction uncertainty (Supplementary Fig. 10).

Variational approximations typically underestimate uncertainty. So, to correct the uncertainty estimates, the model prediction uncertainty obtained from the variational approximation was compared to the model root-mean-square error (RMSE) (i.e. the root-mean-square difference between the model prediction and the experimental measurement). For all three Hill equation parameters (*G*_0_, *G*_∞_, and *EC*_50_), both the prediction uncertainty and the RMSE increase with the number of amino acid substitutions relative to wild-type sequence (Supplementary Fig. 10a-b), and the RMSE at each substitutional distance is an approximately linear function of the median model uncertainty (Supplementary Fig. 10c). So, for the single-substitution analysis (Fig. 2, Fig. 5b-c, Supplementary Fig. 13, Supplementary Data 1), the uncertainties from the variational approximation were multiplied by a factor of 3.8. This rescaled the uncertainties so that the median uncertainty was approximately equal to the RMSE for each substitutional distance.

## Data Availability

The raw sequence data for long-read and short-read DNA sequencing have been deposited in the NCBI Sequence Read Archive and are available under the project accession number PRJNA643436. Plasmid sequences have been deposited in the NCBI Genbank under accession codes MT702633, and MT702634, for pTY1 and pVER, respectively.

The processed data table containing information for each LacI variant in the library is publicly available via the NIST Science Data Portal, with the identifier ark:/88434/mds2-2259 (https://data.nist.gov/od/id/mds2-2259 or https://doi.org/10.18434/M32259).

## Code Availability

All custom data analysis code is available at https://github.com/djross22/nist_lacI_landscape_analysis.

## Acknowledgements

We would like to thank Vanya Paralanov, Daniel Samarov, Ben Scott, Zvi Kelman, Gilad Kusne, and Swarnavo Sarkar for thoughtful discussions during planning and execution of this work. We would also like to thank Jayan Rammohan, William Brad O’Dell, and Elizabeth Strychalski for insights during the experimental work, as well as improving the manuscript.

## Author Contributions

D.S.T., and D.R. conceived of the process.

D.S.T, S.L., and D.R. developed the experimental workflow.

D.S.T. designed, built, and tested genetic constructs. E.F.R., and D.R. programmed automated protocols.

D.S.T., E.F.R., N.A., O.V., and D.R. performed landscape and verification experiments.

P.D.T. and D.R. performed Bayesian inference and model fitting.

P.D.T. designed and evaluated the recurrent architecture for machine learning.

P.D.T., N.D.O, and D.R. contributed to long-read sequencing analysis.

D.S.T., P.D.T, A.P., and D.R. wrote the manuscript. All authors contributed to the manuscript.

## Supplementary information

Supplementary Information: This file includes 17 supplementary figures and 1 supplementary table.

Supplementary Data 1: Single-substitution Hill equation parameters. The table contains the estimated Hill equation parameter for all of the single-substitution LacI variants analyzed. The values listed in the table are the base-10 logarithm for each parameter. The column headings indicate the parameter and the source of the estimate as follows: “exp_”: experimental values, “dnn_”: values predicted by the DNN model, “est_”: values for missing substitutions estimated based on previously published results, “_err”: the uncertainty (1 standard deviation) of the log_10_(parameter). Column headings that start with “best_” contain the values used for the analysis and plots contained in the manuscript. See Methods for more details.

## Disclaimer

The authors declare no competing interests

Certain commercial equipment, instruments, or materials are identified to adequately specify experimental procedures. Such identification neither implies recommendation nor endorsement by the National Institute of Standards and Technology nor that the equipment, instruments, or materials identified are necessarily the best for the purpose.

Supplementary Information is available for this paper.

## Notes

https://doi.org/10.18434/M32259

### Competing Interest Statement

The authors have declared no competing interest.

