## Supplementary Information for "The genotype-phenotype landscape of an allosteric protein"

**This PDF file includes:**

Supplementary Figures 1 to 17
Supplementary Table 1
Caption for Supplementary Data 1

**Other supplementary information for this manuscript include the following:**

Supplementary Data 1, Hill equation parameters for LacI variants with single amino acid substitutions.

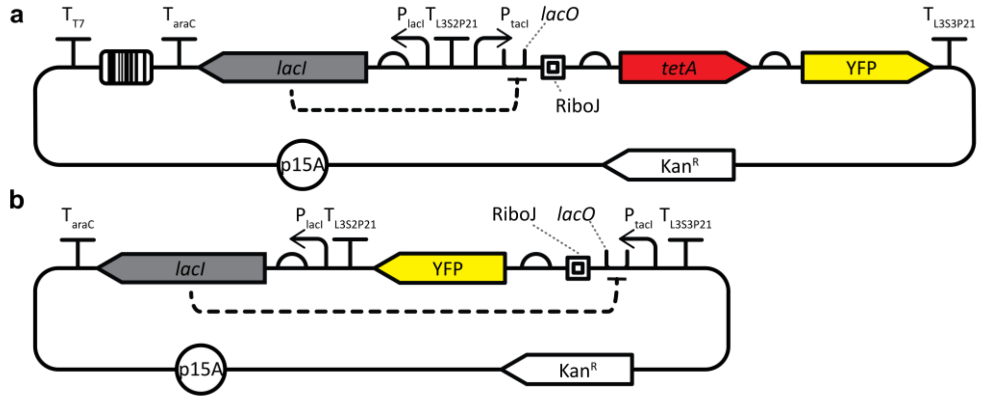


**Supplementary Fig. 1 Plasmid maps for the two plasmids used in this work.** Both plasmids contain the p15A origin of replication (p15A), kanamycin resistance gene ($Kan^R$), and *lacI* coding DNA sequence (*lacI*). In both plasmids, the encoded LacI protein transcriptionally regulates an output that was driven by the $P_{tacI}$ promoter, *lacO* operator, and RiboJ transcriptional insulator. **a** The library plasmid (pTY1) was used for measurement of the genotype and phenotype of the entire LacI library. The LacI library, including the coding DNA sequences of *lacI* variants and their corresponding barcodes, was cloned into pTY1. In pTY1, the LacI variants regulated the expression of a tetracycline resistance gene, *tetA*. Plasmid pTY1 also encoded yellow fluorescence protein (YFP), which was used for cloning purposes. **b** The verification plasmid (pVER) was used to verify the dose-response curves of over 100 variants from the library. To verify the dose-response curve of a LacI variant, the coding DNA sequence for that variant was chemically synthesized and cloned into pVER, where the LacI variant regulated the expression of YFP. The dose-response curve was then measured using flow cytometry.

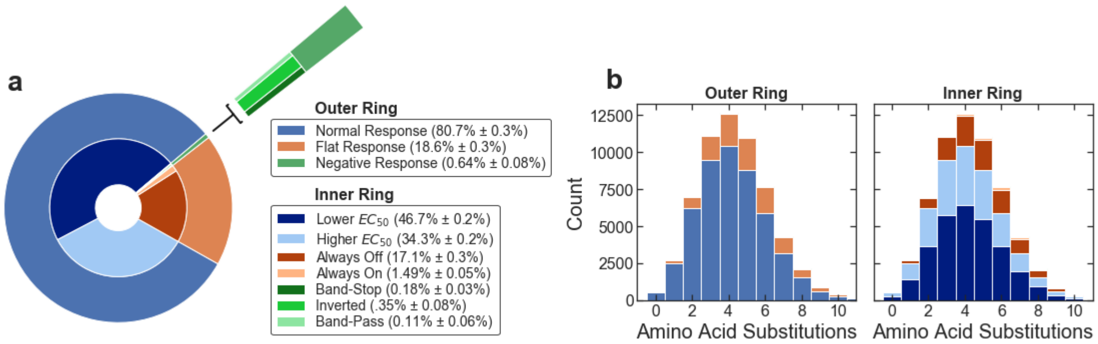


**Supplementary Fig. 2 Diversity of phenotypes in the LacI library. a** Relative abundance of the various LacI phenotypes found in the library. Variants with a "normal response" phenotype have dose-response curves qualitatively similar to the wild-type, with $G_0 < G_\infty$. Normal response variants include variants with $EC_{50}$ lower than the wild-type value (between approximately 1 µmol/L and 100 µmol/L) and variants with $EC_{50}$ higher than the wild-type value (between approximately 100 µmol/L and 2000 µmol/L). Variants with a "flat response" phenotype have flat dose-response curves with $G_0 \cong G_\infty$. Flat response variants include always-off variants ($G(0) < 0.25 \times G_{\infty,wt}$; i.e., the $I^S$ phenotype from Markiewicz *et al*., *Journal of Molecular Biology*, **240**, 421-433, 1994) and always-on variants ($G(0) > 0.25 \times G_{\infty,wt}$; i.e. the $I^-$ phenotype from Markiewicz *et al*.). Variants with a "negative response" phenotype have dose-response curves with a negative slope, $\partial G/\partial L$, for some portion of the measured concentration range. Negative response variants include band-stop, inverted, and band-pass variants. **b** Histogram of the number of variants with each LacI phenotype found in the library as a function of the number of amino acid substitutions. The plots in **a** and **b** share the same legend. The outer ring of **a** and the left panel of **b** show the proportion of variants with dose-response curves that are normal, flat-response, or negative response. The inner ring of **a** and the right panel of **b** show more detailed descriptors of variant phenotypes.

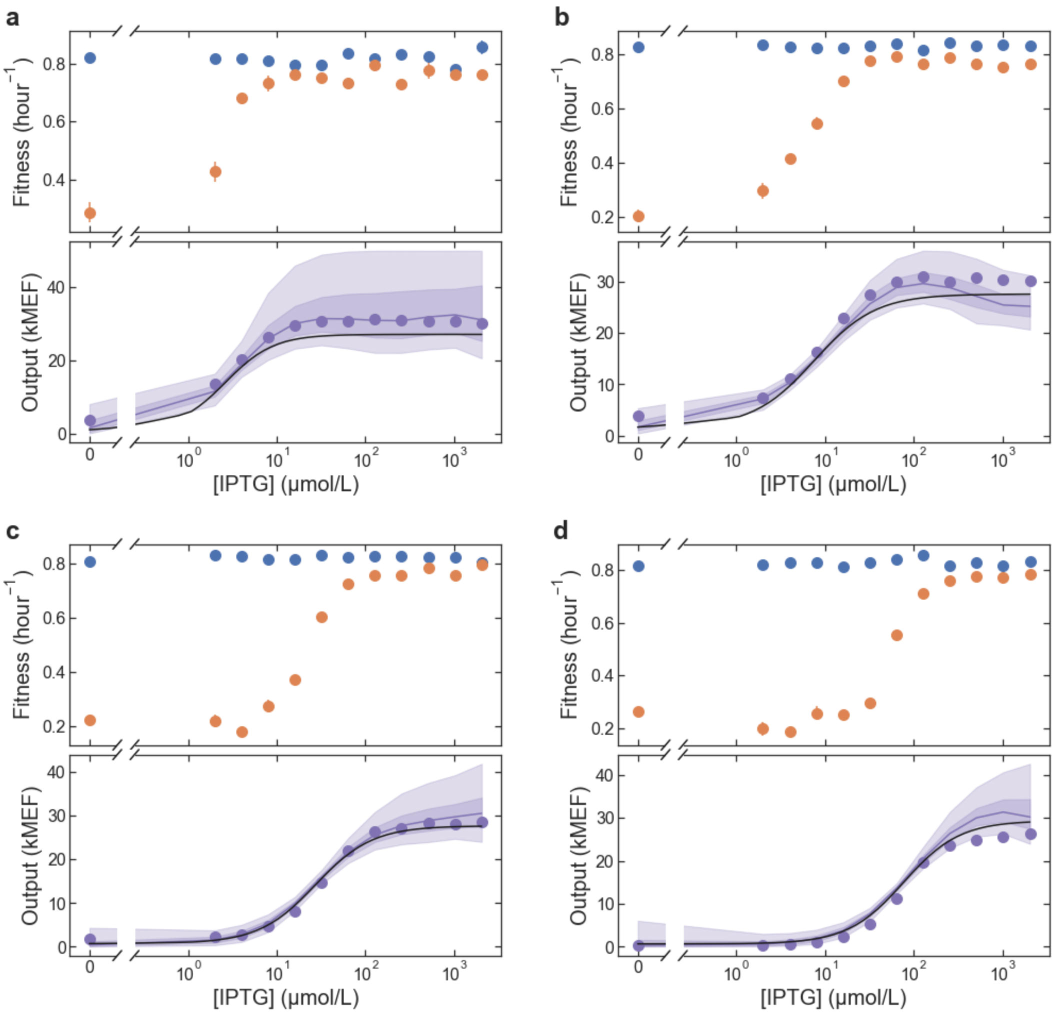


**Supplementary Fig. 3 Example data for fitness and dose-response curves for normal phenotype LacI variants. a-d** Each pair of plots shows the fitness (top) and dose-response curve (bottom) for a different LacI variant. The fitness data is from the library-scale landscape measurement. The fitness without tetracycline is shown as blue points and the fitness with tetracycline is shown as orange points. In the dose-response plots, the black lines show the results from the library-scale measurement using the Bayesian Hill equation model. The purple lines and shaded regions show the results from the library-scale measurement using the Bayesian Gaussian process (GP) model, where the line is the median GP results and the shaded regions indicate 50% and 90% credible intervals. The plotted purple points show the dose-response curves from the flow cytometry verification measurements. Error bars indicate ± one standard deviation and are often within the markers.

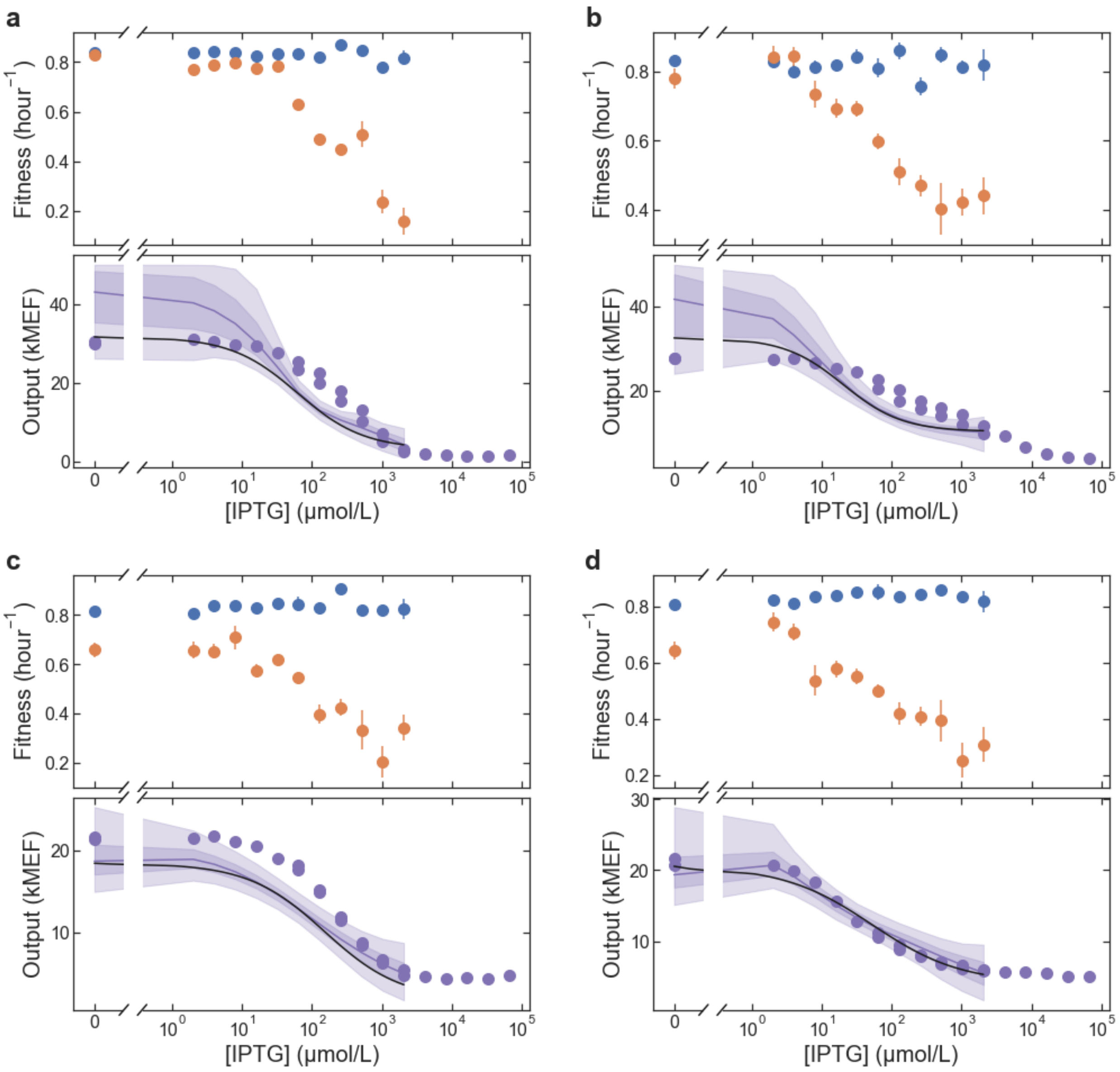


**Supplementary Fig. 4 Example data for fitness and dose-response curves for inverted phenotype LacI variants. a-d** Each pair of plots shows the fitness (top) and dose-response curve (bottom) for a different LacI variant. The fitness data is from the library-scale landscape measurement. The fitness without tetracycline is shown as blue points and the fitness with tetracycline is shown as orange points. In the dose-response plots, the black lines show the results from the library-scale measurement using the Bayesian Hill equation model. The purple lines and shaded regions show the results from the library-scale measurement using the Bayesian Gaussian process (GP) model, where the line is the median GP results and the shaded regions indicate 50% and 90% credible intervals. The plotted purple points show the dose-response curves from the flow cytometry verification measurements. Error bars indicate ± one standard deviation and are often within the markers.

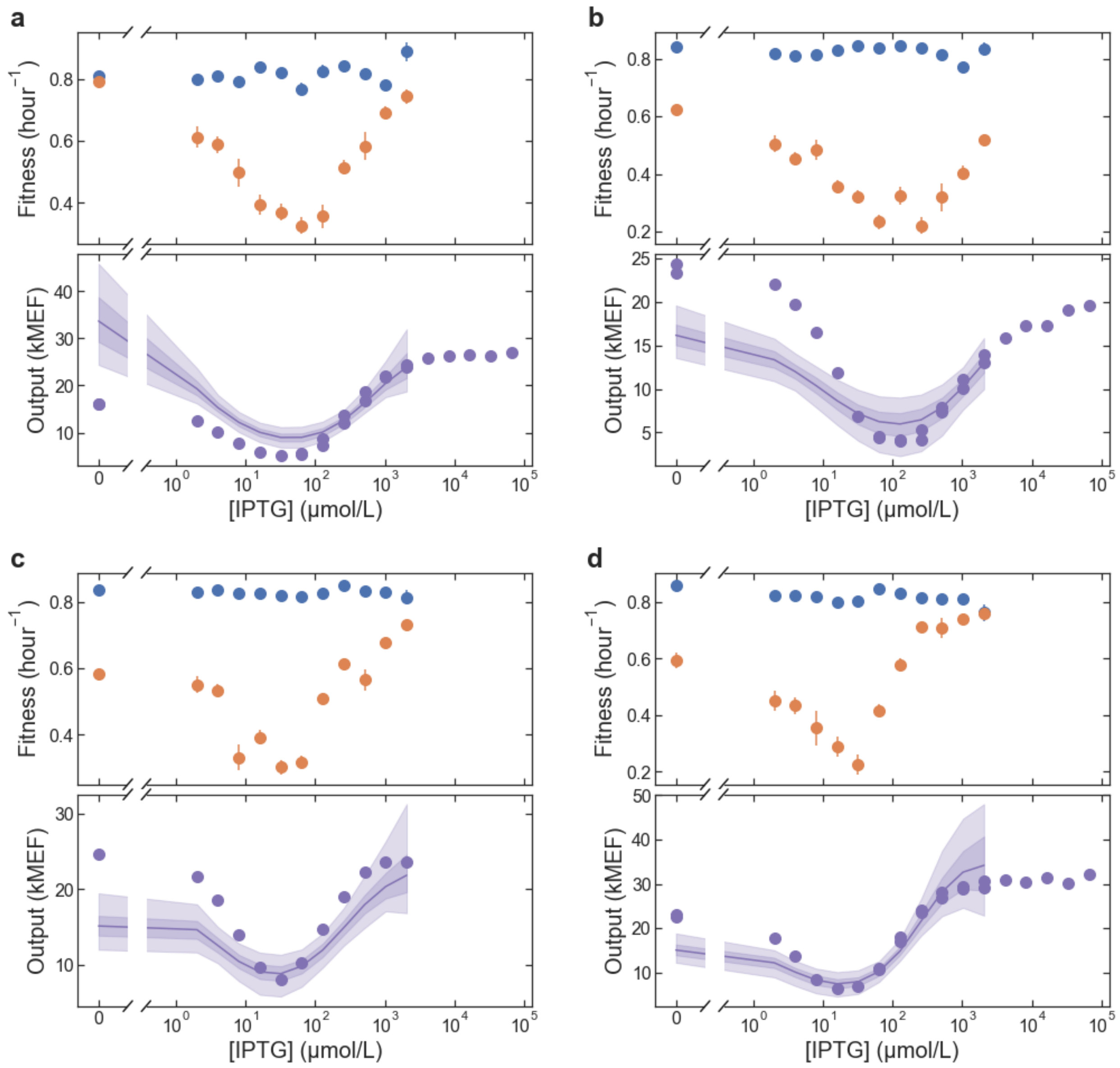


**Supplementary Fig. Example data for fitness and dose-response curves for band-stop phenotype LacI variants. a-d** Each pair of plots shows the fitness (top) and dose-response curve (bottom) for a different LacI variant. The fitness data is from the library-scale landscape measurement. The fitness without tetracycline is shown as blue points and the fitness with tetracycline is shown as orange points. In the dose-response plots, the black lines show the results from the library-scale measurement using the Bayesian Hill equation model. The purple lines and shaded regions show the results from the library-scale measurement using the Bayesian Gaussian process (GP) model, where the line is the median GP results and the shaded regions indicate 50% and 90% credible intervals. The plotted purple points show the dose-response curves from the flow cytometry verification measurements. Error bars indicate ± one standard deviation and are often within the markers.

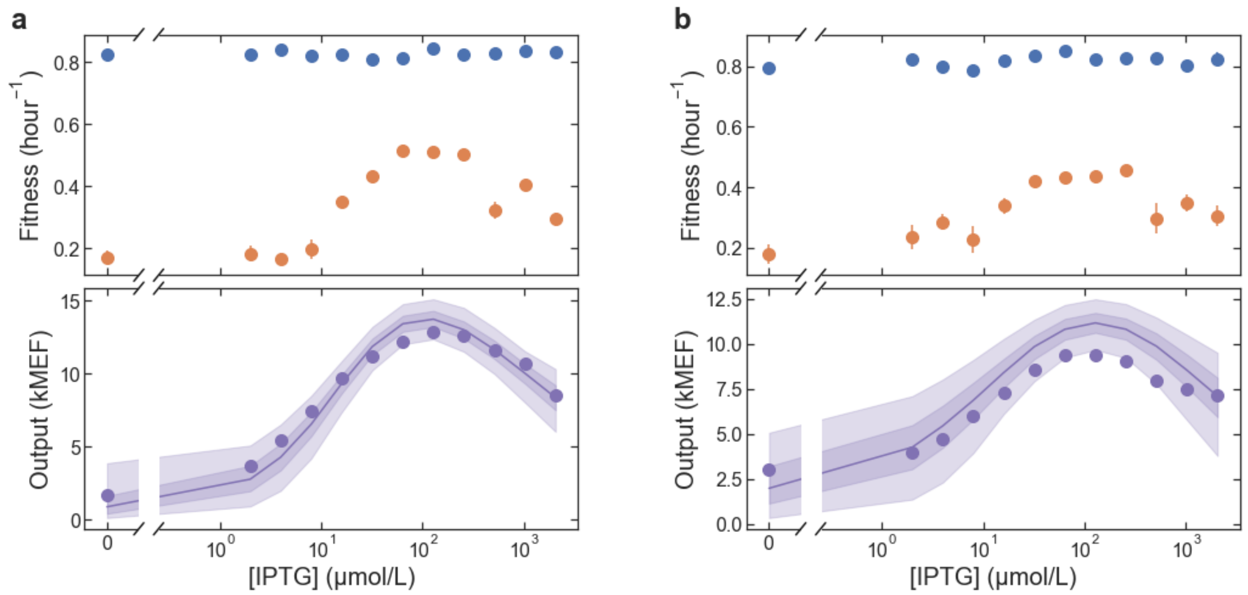


**Supplementary Fig. 6 Example data for fitness and dose-response curves for band-pass phenotype LacI variants. a-d** Each pair of plots shows the fitness (top) and dose-response curve (bottom) for a different LacI variant. The fitness data is from the library-scale landscape measurement. The fitness without tetracycline is shown as blue points and the fitness with tetracycline is shown as orange points. In the dose-response plots, the black lines show the results from the library-scale measurement using the Bayesian Hill equation model. The purple lines and shaded regions show the results from the library-scale measurement using the Bayesian Gaussian process (GP) model, where the line is the median GP results and the shaded regions indicate 50% and 90% credible intervals. The plotted purple points show the dose-response curves from the flow cytometry verification measurements. Error bars indicate ± one standard deviation and are often within the markers.

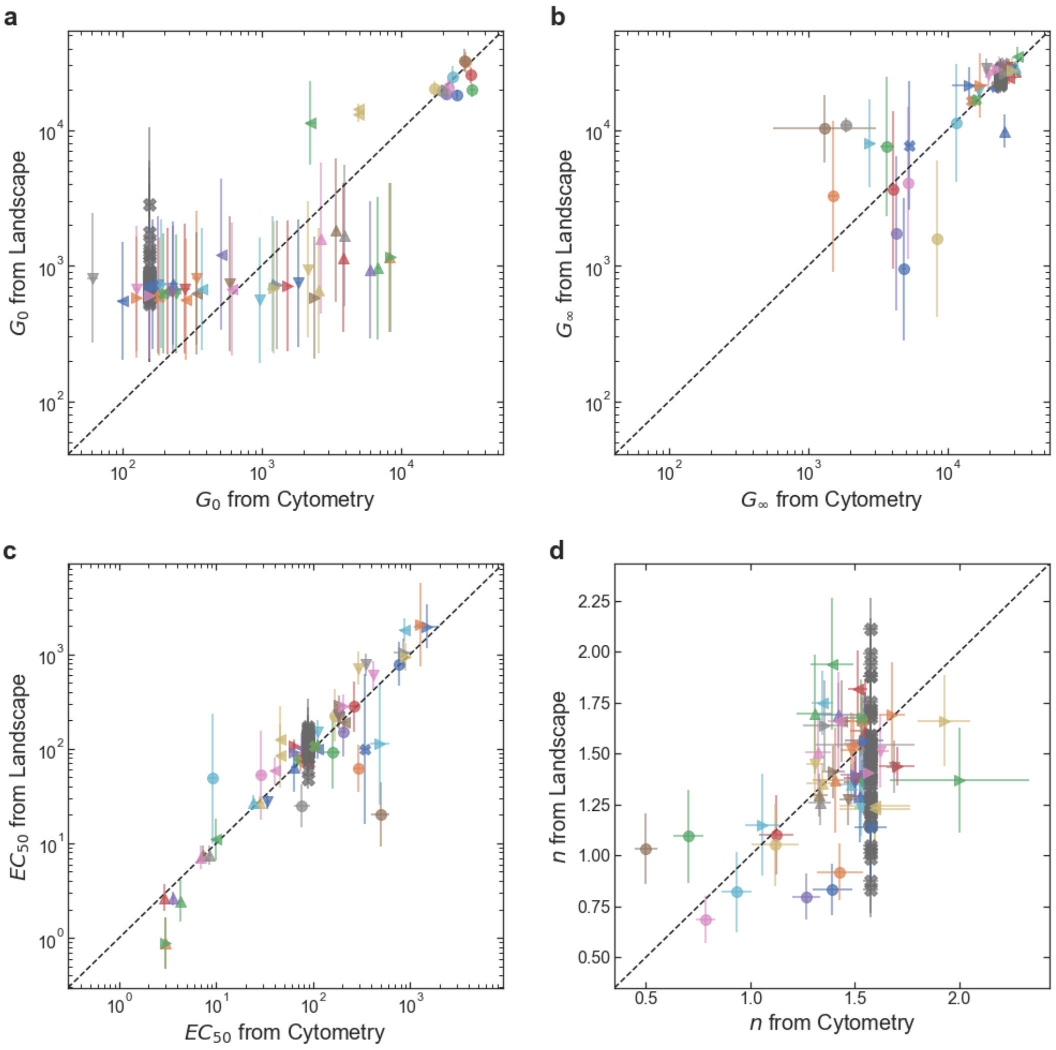


**Supplementary Fig. 7 Accuracy of the library-scale dose-response curve measurement. a-d** The plots compare the results from the library-scale measurement (y-axis) with the flow cytometry verification results (x-axis) for each Hill equation parameter. Data is shown for all of the verified LacI variants with sigmoidal dose-response curves (i.e. band-stop and band-pass variants are not included). Data for different variants are plotted with different combinations of color and shape. Variants that occurred more than once in the library (with different DNA barcodes) are plotted multiple times. For example, the wild type (dark gray 'X' symbols) is plotted 53 times. The accuracy for each Hill equation parameter is: 4-fold for $G_0$ (**a**), 1.5-fold for $G_\infty$ (**b**), 1.8-fold for $EC_{50}$ (**c**), and ± 0.28 for $n$ (**d**). For $G_0$, $G_\infty$, and $EC_{50}$ (**a-c**), the accuracy is calculated as: $\exp\left(\mathrm{RMSE}\left(\ln(x)\right)\right)$, where $\mathrm{RMSE}\left(\ln(x)\right)$ is the root-mean-square difference between the logarithm of each parameter from the library-scale and cytometry measurements. For $n$, the accuracy is given simply as the root-mean-square difference between the library-scale and cytometry results. Error bars indicate ± one standard deviation.

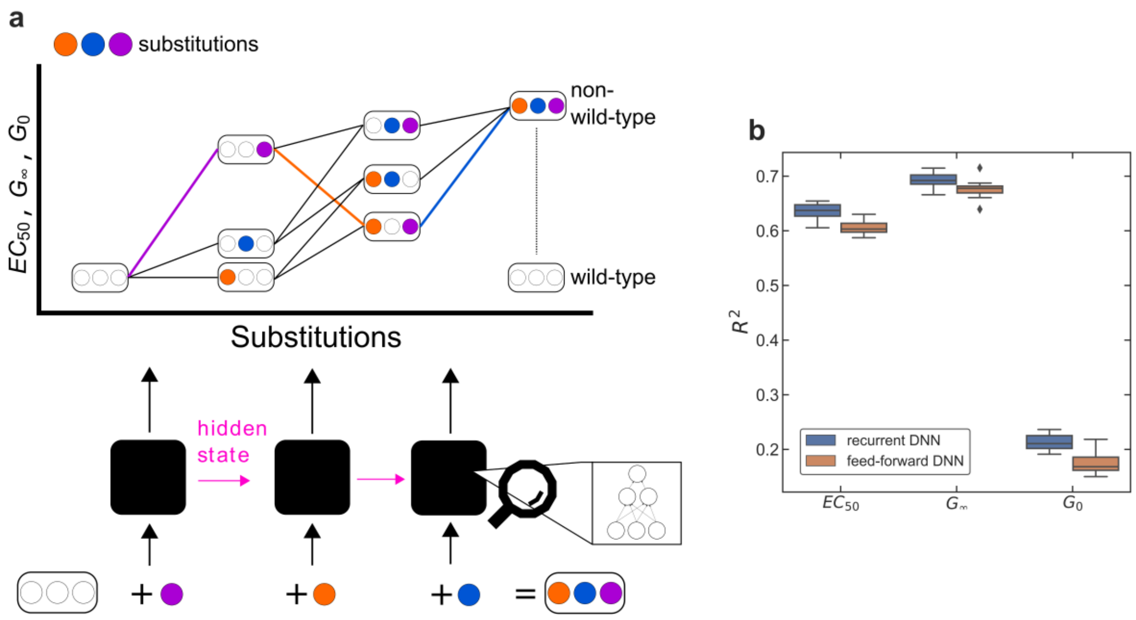


**Supplementary Fig. 8 Deep neural network (DNN) model. a** Schematic illustration of recurrent deep neural network model architecture. Using the wild-type sequence as a starting point, the model predicts the Hill equation parameters for non-wild-type sequences by composing the individual parameter changes due to sequential amino acid substitutions along a mutational path. These changes are then added together to predict the final parameter values. All paths to a non-wild-type sequence converge to the same value, and this fact is leveraged to build a recurrent neural network model that learns to predict the effects of each substitution, given all previous substitutions. Potential non-additive effects are captured by the hidden state of the model, which predicts the change in parameter value for the most recent substitution and serves as a set of latent variables for predicting subsequent substitutions. Note that variants with intermediate sequences may be present in the library, but this is not necessary to train the model. The model will still learn to predict intermediate steps in the path, even if that data is not present. **b** Performance of recurrent and feed-forward DNN models. The boxplot summarizes the $R^2$ values for ten cross-validation tests for each model. The recurrent DNN model generally outperforms the feed-forward model, giving higher $R^2$ values for each of the Hill equation parameters.

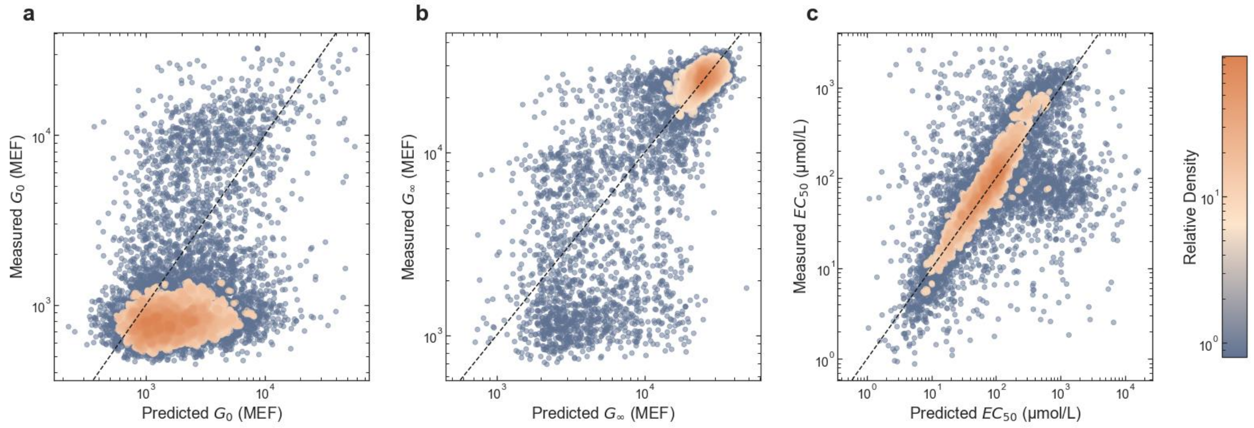


**Supplementary Fig. 9 Prediction of LacI variant phenotypes with recurrent deep neural network (DNN) model. a-c** Measured vs. predicted values for each Hill equation parameter. The plotted results only show holdout data not used in model training. Predictions are taken from the variational posterior mean for each Hill equation parameter. Measured values are the posterior medians from the Bayesian fits to the experimental data. In **a-b**, $G_\infty$ is given in molecules of equivalent fluorophore (MEF). In **c**, the cluster of points with measured $EC_{50}$ near 100 µmol/L and predicted $EC_{50}$ near 1000 µmol/L correspond to variants for which the DNN model provided better $EC_{50}$ and $G_\infty$ estimates than the measurement, because $EC_{50}$ was near or above the maximum ligand concentration used. For these points, the nearly constant dose-response across the measured concentration range resulted in a large uncertainty for $EC_{50}$ and a posterior median near the median of the $EC_{50}$ prior (100 µmol/L).

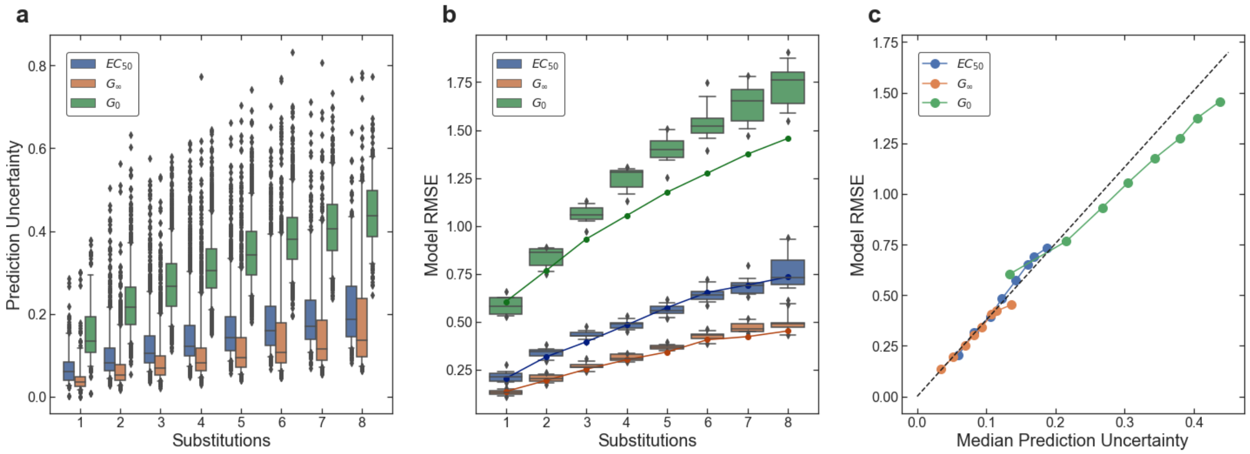


**Supplementary Fig. 10 DNN model prediction uncertainty and root-mean-square error (RMSE).**
**a** Model prediction uncertainty for the base-ten logarithm of each Hill equation parameter as a function of the number of amino acid substitutions relative to the wild-type LacI sequence. The boxplot shows the distribution of posterior standard deviation values for the set of variants with the indicated number of substitutions. **b** Model RMSE for each Hill equation parameter as a function of the number of substitutions relative to the wild-type sequence. RMSE is the root-mean-square difference between the model prediction and experimental measurement for the base-ten logarithm of each Hill equation parameter. The boxplot shows the distribution of RMSE values from the ten cross-validation tests for the recurrent DNN model. Solid lines with points show the RMSE for the holdout data not used for training. Both the prediction uncertainty (**a**) and the RMSE (**b**) increase with increasing number of substitutions relative to the wild-type sequence. **c** Model RMSE as a function of the median prediction uncertainty for each Hill equation parameter. Each plotted point is for a different number of substitutions. The RMSE values are from the holdout data, i.e. the solid lines with points from (**b**). The median prediction accuracy values are from the distributions plotted in (**a**). The dashed line indicates the multiplicative factor (approximately 3.8) used to correct the uncertainty values (see Methods).

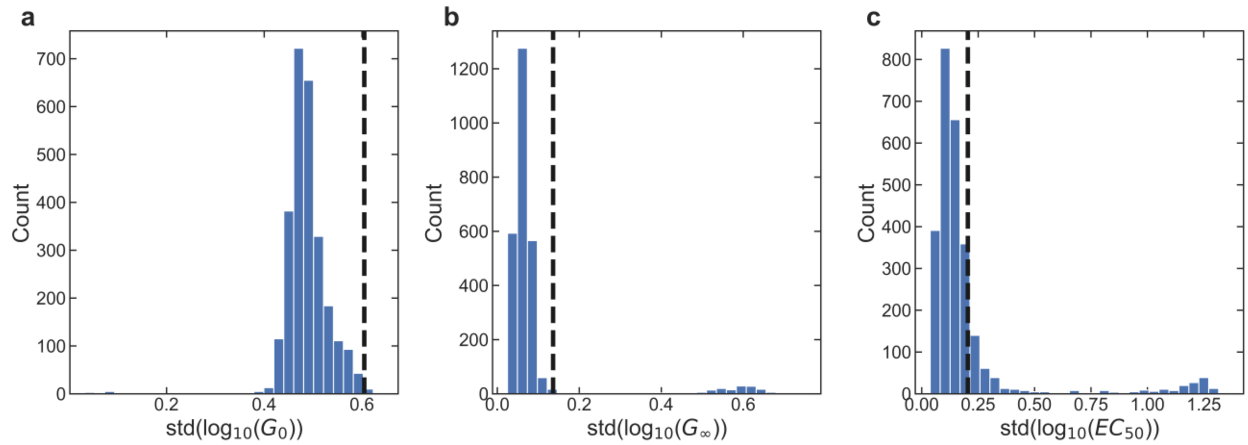


**Supplementary Fig. 11 Comparison between the DNN model root-mean-square error (RMSE) and the measurement uncertainty for single mutants. a-c** The histograms show distributions of the measurement uncertainty for LacI variants with single amino acid substitutions (relative to the wild type) for each Hill equation parameter. For comparison, the dashed line in each plot shows the RMSE for the DNN model predictions for single-substitution hold-out data. The model RMSE as a function of the number of substitutions is shown in Supplementary Fig. 10b.

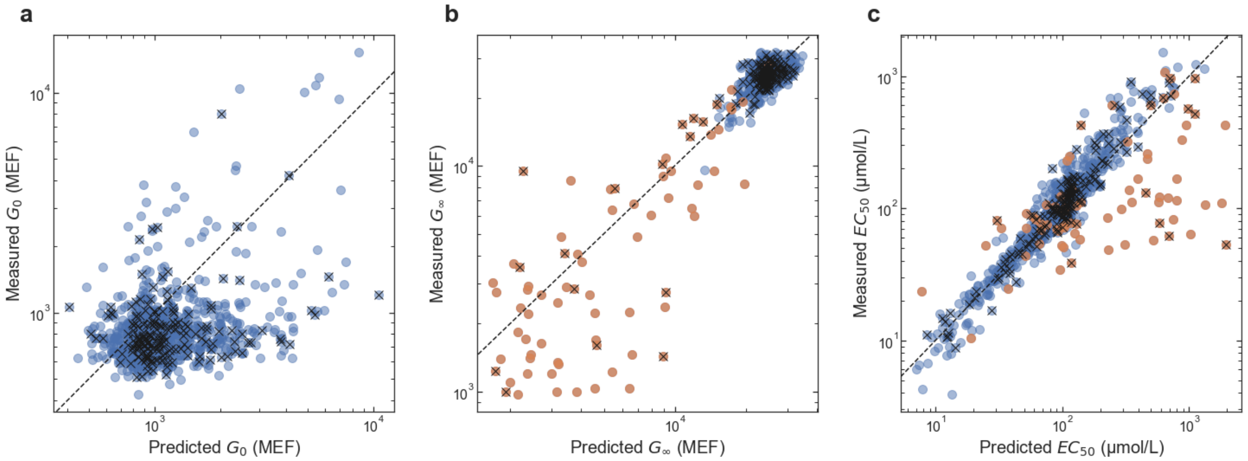


**Supplementary Fig. 12 Measured vs. predicted values from DNN model for single amino acid substitutions**. **a-c** The plots compare the DNN model predictions (x-axis) with the measured values (y-axis) for the Hill equation parameters for LacI variants with single amino acid substitutions. Blue symbols show data for all of the single-substitution variants in the library. Orange symbols show data for variants with a high uncertainty for the measured $EC_{50}$ (std($\log_{10}(EC_{50})$) > 0.35). The $EC_{50}$ uncertainty for those variants was relatively high either because $G_\infty$ was similar to $G_0$ and/or because $EC_{50}$ was near or above the maximum ligand concentration used (2048 µmol/L). For those variants, in the analysis of single-mutant effects, the DNN model result for $G_\infty$ and $EC_{50}$ was used in place of the experimental result. Points marked with an 'x' were in the holdout data not used for model training. In all plots, $G_0$ and $G_\infty$ are given in molecules of equivalent fluorophore (MEF).

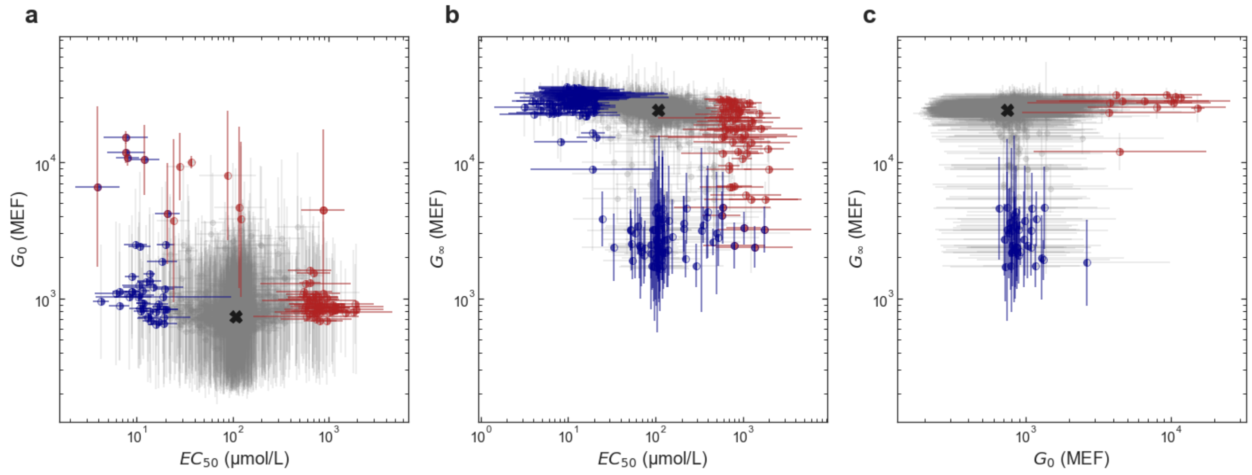


**Supplementary Fig. 13 Multiparametric impact of amino acid substitutions on allosteric function.**
**a-c** Each plot shows the joint effect of single amino acid substitutions on two Hill equation parameters. In each plot, substitutions that change both Hill equation parameters by less than 5-fold are shown as light gray points, and substitutions that change one or both Hill equation parameters by more than 5-fold are shown as red or blue points with error bars. As in Fig. 2 (in the main manuscript), red indicates a decrease in Hill equation parameter and blue indicates an increase. The left half of each symbol and the y-error bar are colored based on the y-axis parameter; the right half of each symbol and the x-error bars are colored based on the x-axis parameter. The wild-type phenotype is indicated with a black 'X' in each plot. In all plots, $G_0$ and $G_\infty$ are given in molecules of equivalent fluorophore (MEF). Error bars indicate ± one standard deviation.

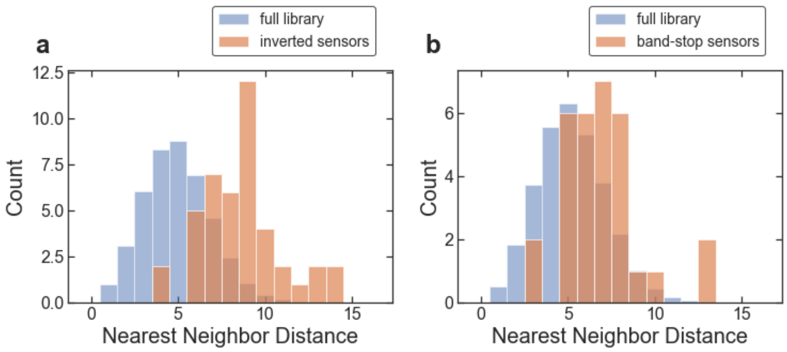


**Supplementary Fig. 14 Nearest neighbor distance histograms for inverted and band-stop genotypes**. **a-b** In each plot, the orange bars show the distribution of nearest neighbor Hamming distance for the amino acid sequences for strongly inverted (**a**) and strong band-stop (**b**) variants, and the blue bars show the distribution of nearest neighbor Hamming distance for a similar number of randomly selected sequences from the full library. The full-library histograms (blue bars) are averaged over 1000 iterations of randomly selected sequences.

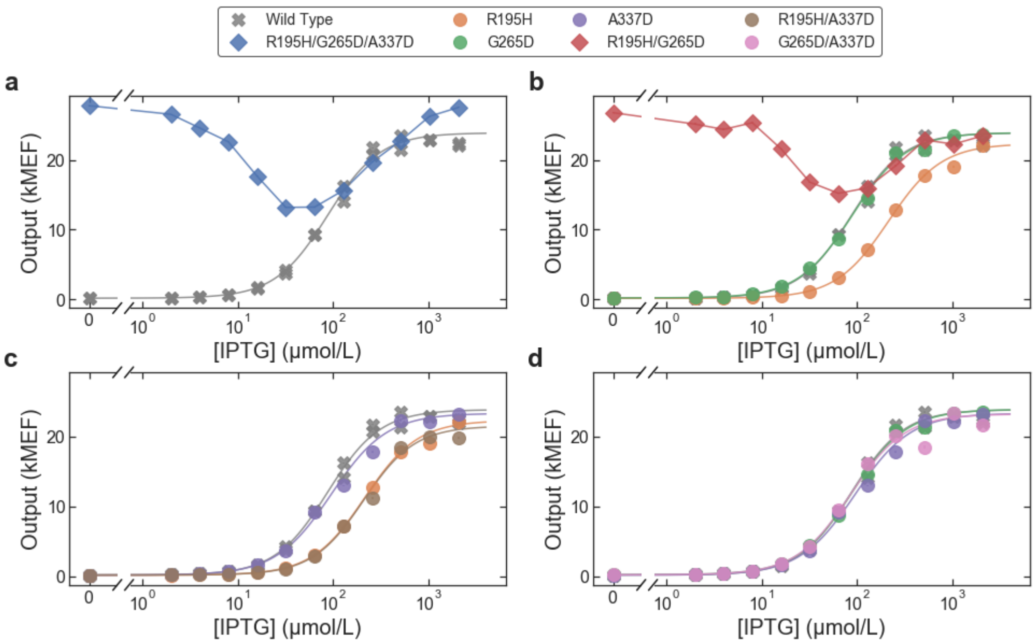


**Supplementary Fig. 15 The band-stop phenotype emerges from the combination of two amino acid substitutions. a** Dose-response curves measured with flow cytometry for wild-type LacI and a strong band-stop variant identified from the library with only three amino acid substitutions (R195H, G265D, A337D). **b-d** Dose-response curves for LacI variants containing single- and double-substitution permutations of the three substitutions in the band-stop variant. Each plot shows two LacI variants containing single substitutions and one LacI variant containing both substitutions. The single substitutions R195H (orange) or G265D (green) result in sigmoidal dose-response curves similar to the wild-type, but the combination of the two, R195H/G265D (red), results in a band-stop phenotype (**b**). All other combinations of single- and double-substitutions result in sigmoidal dose-response curves similar to the wild-type (**c-d**).

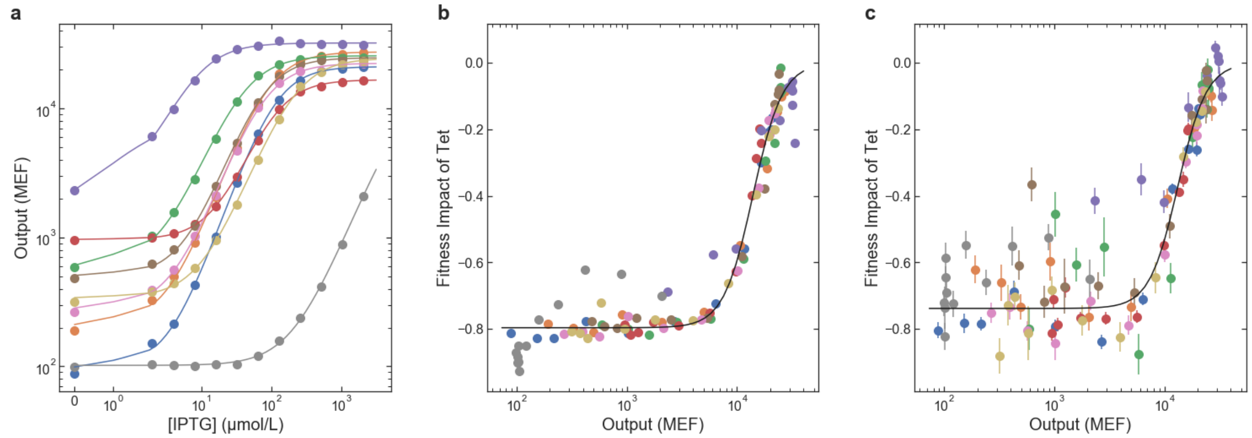


**Supplementary Fig. 16 Calibration data for the determination of dose-response curves using barcode sequencing. a** Dose-response curves measured with flow cytometry for nine LacI variants used to calibrate library-scale measurement. **b-c** The fitness impact of tetracycline (from the library-scale measurement) plotted vs. gene expression output (from flow cytometry of individual variants). The fitness impact of tetracycline is defined as the decrease in fitness (*E. coli* growth rate) measured with tetracycline vs. without tetracycline normalized by the fitness measured without tetracycline, i.e. ($\mu^{tet}/\mu^{0} - 1$) from Eq. (9) in the Methods. **b** Data from a small-scale test library containing only the calibration variants. **c** Data from measurement of the full library. Results for different calibration variants are shown with different colored symbols. The solid black lines in **b-c** show the results of a fit using Eq. (9). Error bars indicate ± one standard deviation. The results for the full library measurement are noisier than the small-scale test because of the lower read count per variant (approximately 100-fold difference), but the general trend and the fits for both results are similar.

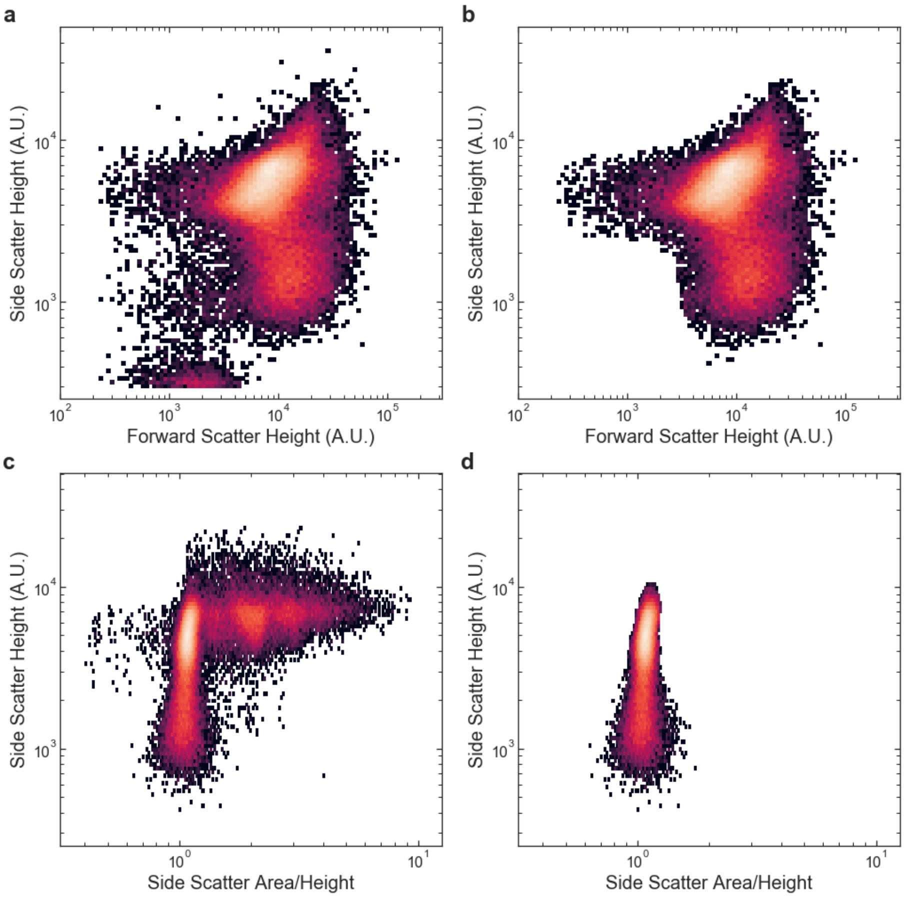


**Supplementary Fig. 17 Flow cytometry gating example. a** Side scatter vs. forward scatter plot before automated cell gating, showing both cell and non-cell detection events. **b** Side scatter vs. forward scatter plot after automated cell gating, showing only events most likely to be cell events. **c** Side scatter vs. side scatter area/height plot before automated singlet gating, showing both singlet and multiplet cell detection events. **d** Side scatter vs. side scatter area/height plot after automated singlet gating, showing only singlet cell detection events.

| region | start | first 12 bases | end | last 12 bases | nominal length | mutation rate |
|---|---|---|---|---|---|---|
| barcode 1 | 116 | NNTNNNANNTNN | 152 | NNANNTNNNANN | 37 | N/A |
| inter-barcode | 153 | ATATGCCAGCAG | 176 | GCCGGCCACGCT | 24 | 0.132 |
| barcode 2 | 177 | NNTNNNANNTNN | 213 | NNANNTNNNANN | 37 | N/A |
| intergenic | 214 | CGGTGGCCCGGG | 394 | CCTCCTGGATTA | 181 | 0.352 |
| lacI | 395 | TCACTGCCCGCT | 1477 | TACTGGTTTCAT | 1083 | 6.478 |
| regulatory | 1478 | ATTCACCACCCT | 1789 | GAGAGCTGCTAC | 312 | 0.171 |
| tetA | 1790 | ATGAGTAGCAGT | 2992 | GAAACGAGTGCC | 1203 | 0.018 |
| YFP | 2993 | TAACGGCGTAAG | 3735 | CTGTATAAATAA | 743 | 0.021 |
| KAN | 3736 | AAGCGGGAGACC | 4779 | GCGTCAGACCCC | 1044 | 0.008 |
| origin of replication | 4780 | TTAATAAGATGA | 5584 | TGCCAACATAGT | 805 | 0.012 |
| origin + intergenic | 5585 | AAGCCAGTATAC | 115 | GGCTGTCGGCGT | 192 | 0.181 |

**Supplementary Table 1 Regions extracted from long-read sequencing data for library-scale analysis.** The lacI region includes the *lacI* coding DNA sequence (CDS) and stop codon. The regulatory region includes the $P_{lacI}$ and $P_{tacI}$ promoters, the *lacO* operator, the riboJ insulator, and the RBS sites for both *lacI* and *tetA*. The tetA region includes the tetA CDS. The YFP region includes the YFP CDS and its RBS. The KAN region includes the transcriptional promoter and CDS for the kanamycin resistance gene as well as the transcriptional terminator for the genes regulated by LacI ($T_{L3S3P21}$). The origin of replication was split by the start of the PacBio circular consensus HiFi reads, so the last 32 bases of the origin of replication were grouped with the intergenic region between the origin and the barcodes. The start and end columns in the table give the nucleotide position of the start and end of each region based on the nominal full plasmid (GenBank ID: MT702633). The mutation rate column lists the estimated mean number of single nucleotide changes per 1000 bases for each region, using the nominal/wild-type sequence as the reference, and considers only single nucleotide polymorphisms (i.e. no indels). Regions adjacent to the barcodes and the lacI region have elevated mutation rates, due to the methods used for library generation and assembly.

**Supplementary Data 1 Hill equation parameters for LacI variants with single amino acid substitutions (separate file).**

The .csv file contains a table with the estimated Hill equation parameters from both experimental measurements and DNN predictions for all of the single-substitution LacI variants analyzed. The values listed in the table are the base-10 logarithm for each parameter. The column headings indicate the parameter and the source of the estimate as follows: "exp_": experimental values, "dnn_": values predicted by the DNN model, "est_": values for missing substitutions estimated based on previously published results, "_err": the uncertainty (1 standard deviation) of the $\log_{10}$(parameter). Column headings that start with "best_" contain the values used for the analysis and plots contained in the manuscript. See Methods for more details.
